# Activation of oral epithelial EphA2-EFGR signaling by *Candida albicans* virulence factors

**DOI:** 10.1101/491076

**Authors:** Marc Swidergall, Norma V. Solis, Nicolas Millet, Manning Y. Huang, Jianfeng Lin, Quynh T. Phan, Michael D. Lazarus, Zeping Wang, Aaron P. Mitchell, Scott G. Filler

## Abstract

During oropharyngeal candidiasis (OPC), *Candida albicans* invades and damages oral epithelial cells, which respond by producing proinflammatory mediators that recruit phagocytes to foci of infection. The ephrin type-A receptor 2 (EphA2) detects β-glucan and plays a central role in stimulating epithelial cells to release proinflammatory mediators during OPC. The epidermal growth factor receptor (EGFR) also interacts with *C. albicans* and is known to be activated by the Als3 adhesin/invasin and the Ece1/Candidalysin pore-forming toxin. Here, we investigated the interactions among EphA2, EGFR, Als3 and Ece1/Candidalysin during OPC. We found that Als3 and Ece1/Candidalysin function in the same pathway to damage epithelial cells *in vitro*. They also work together to cause OPC in mice. EGFR and EphA2 constitutively associate with each other as part of a physical complex and are mutually dependent for *C. albicans-induced* activation. *In vitro*, either Als3 or Ece1/Candidalysin is required for *C. albicans* to activate EGFR, sustain EphA2 activation, and stimulate epithelial cells to secrete CXCL8/IL-8 and CCL20. In the mouse model of OPC, Ece1/Candidalysin alone activates EGFR and induces CXCL1/KC and CCL20 production. Ece1/Candidalysin is also necessary for the production of IL-1α and IL-17A independently of Als3 and EGFR. These results delineate the complex interplay between host cell receptors and *C. albicans* virulence factors during the induction of OPC and the resulting oral inflammatory response.

**Author summary:** Oropharyngeal candidiasis occurs when the fungus *Candida albicans* proliferates in the mouth. The disease is characterized by fungal invasion of the superficial epithelium and a localized inflammatory response. Two *C. albicans* virulence factors contribute to the pathogenesis of OPC, Als3 which enables the organisms to adhere to and invade host cells and Ece1/Candidalysin which is pore-forming toxin that damages host cells. Two epithelial cell receptors, ephrin type-A receptor 2 (EphA2) and the epidermal growth factor receptor (EGFR) are activated by *C. albicans*. Here, we show that EphA2 and EGFR form part of complex and that each receptor is required to activate the other. Als3 and Ece1/Candidalysin function in the same pathway to damage epithelial cells. In isolated epithelial cells, both of these virulence factors activate EphA2 and EGFR, and stimulate the production of inflammatory mediators. In the mouse model of OPC, Ece1/Candidalysin elicits of a subset of the oral inflammatory response. Of the cytokines and chemokines induced by this toxin, some require the activation of EGFR while others are induced independently of EGFR. This work provides a deeper understanding of the interactions among *C. albicans* virulence factors and host cell receptors during OPC.

## Introduction

Oropharyngeal candidiasis is characterized by superficial fungal invasion of the oral mucosa, epithelial cell death, and leukocyte recruitment to the focus of infection [1]. *C. albicans* can invade the epithelial cell lining of the oropharynx by two mechanisms, active penetration and induced endocytosis. Active penetration occurs when a progressively elongating hypha physically pushes its way into the epithelial cell [2]. Induced endocytosis occurs when invasins such as Als1, Als3 and Ssa1 expressed by a hypha interact with epithelial cell receptors such as E-cadherin, HER2, and the epidermal growth factor receptor (EGFR), stimulating the epithelial cell to endocytose the organism [3–5]. *C. albicans* also damages epithelial cells by secreting Ece1, which is processed by the Kex2 protease to form Candidalysin, a pore-forming toxin [6, 7].

The epithelial cells that line the oropharynx sense the presence of *C. albicans* and orchestrate the host inflammatory response to fungal overgrowth. In addition to producing host defense peptides that have direct antifungal activity, oral epithelial cells secrete alarmins, proinflammatory cytokines, and chemokines that recruit phagocytes to foci of infection and enhance their candidacidal activity to limit the growth of the invading fungus [8–11]. This epithelial cell response is amplified by interleukin (IL)-17, which is secreted by γδ T cells, innate TCRαβ^+^ cells, and type-3 innate lymphoid cells [12–15].

Recently, it has become clear that the proinflammatory response to *C. albicans* is triggered when the fungus activates specific epithelial cell receptors. We determined that the ephrin type-A receptor 2 (EphA2) is an epithelial cell receptor that senses exposed β-glucan on the fungal surface. When *C. albicans* proliferates, EphA2 is activated, stimulating oral epithelial cells to secrete host defense peptides and proinflammatory mediators. In mice, EphA2 activation is required for the early production of IL-17A, and *Epha2*^-/-^ mice are highly susceptible to the initial stages of OPC [16, 17]. Although EphA2 is required for the normal host defense against OPC, exposure of oral epithelial cells to purified β-glucan *in vitro* induces only transient EphA2 activation and is not sufficient to initiate a significant inflammatory response. By contrast, exposure to live *C. albicans* induces sustained EphA2 activation and a strong inflammatory response [16].

Another proinflammatory epithelial cell stimulus is Candidalysin. This toxin causes an influx of calcium into the epithelial cells, which activates matrix metalloproteases and induces shedding of native EGFR ligands, leading to activation of EGFR signaling and a proinflammatory response by epithelial cells [18].

In the current study, we sought to elucidate how *C. albicans* infection prolongs EphA2 activation and induces a proinflammatory response during OPC. We found that *in vitro* Als3 and Ece1 function in the same pathway to activate epithelial cell EGFR, which maintains EphA2 phosphorylation and stimulates the secretion of CXCL8/IL-8 and CCL20. In mice with acute OPC, pharmacological inhibition of EGFR leads to a reduction in oral fungal burden and impaired inflammatory response during disease progression. Maximal mucosal infection requires Als1 and Als3, which act in the same pathway as Ece1, while the mucosal inflammatory response is largely driven by Ece1. Thus, *C. albicans* activation of EGFR mediates both fungal invasion of the epithelium and induction of the local inflammatory responses to this fungus.

## Results

### The cellular fate of epithelial EphA2 differs depending on the type of stimulation

When oral epithelial cells are infected with yeast-phase *C. albicans*, the fungus germinates and begins to form hyphae within 60 min. Previously, we found that *C. albicans* infection induces EphA2 autophosphorylation on serine 897 within 15 min of infection and that this phosphorylation is sustained for at least 90 min. By contrast, when oral epithelial cells are exposed to purified β-glucan in the form of zymosan or laminarin, EphA2 is phosphorylated for only the first 30 min of contact. At later time points, EphA2 phosphorylation returns to basal levels, even though β-glucan is still present [16]. To investigate how oral epithelial cells respond to prolonged exposure to different EphA2 agonists, we compared their response to the native EphA2 ligand, ephrin A1(EFNA1) and to *C. albicans*. After 15 min of exposure, both stimuli induced phosphorylation of EphA2 (Fig 1A, S1A Fig). After 60 min of exposure, the two stimuli induced distinctly different responses. In cells exposed to EFNA1, there was minimal phosphorylation of EphA2, and total EphA2 levels declined (Fig 1B, S1B and S1C Fig). In cells exposed to *C. albicans*, EphA2 phosphorylation was sustained and there was no decrease in total EphA2 levels. Thus, exposure to *C. albicans* not only induces EphA2 phosphorylation, but prevents the subsequent EphA2 degradation that normally occurs with prolonged exposure to EFNA1 [19].

**Fig 1.**
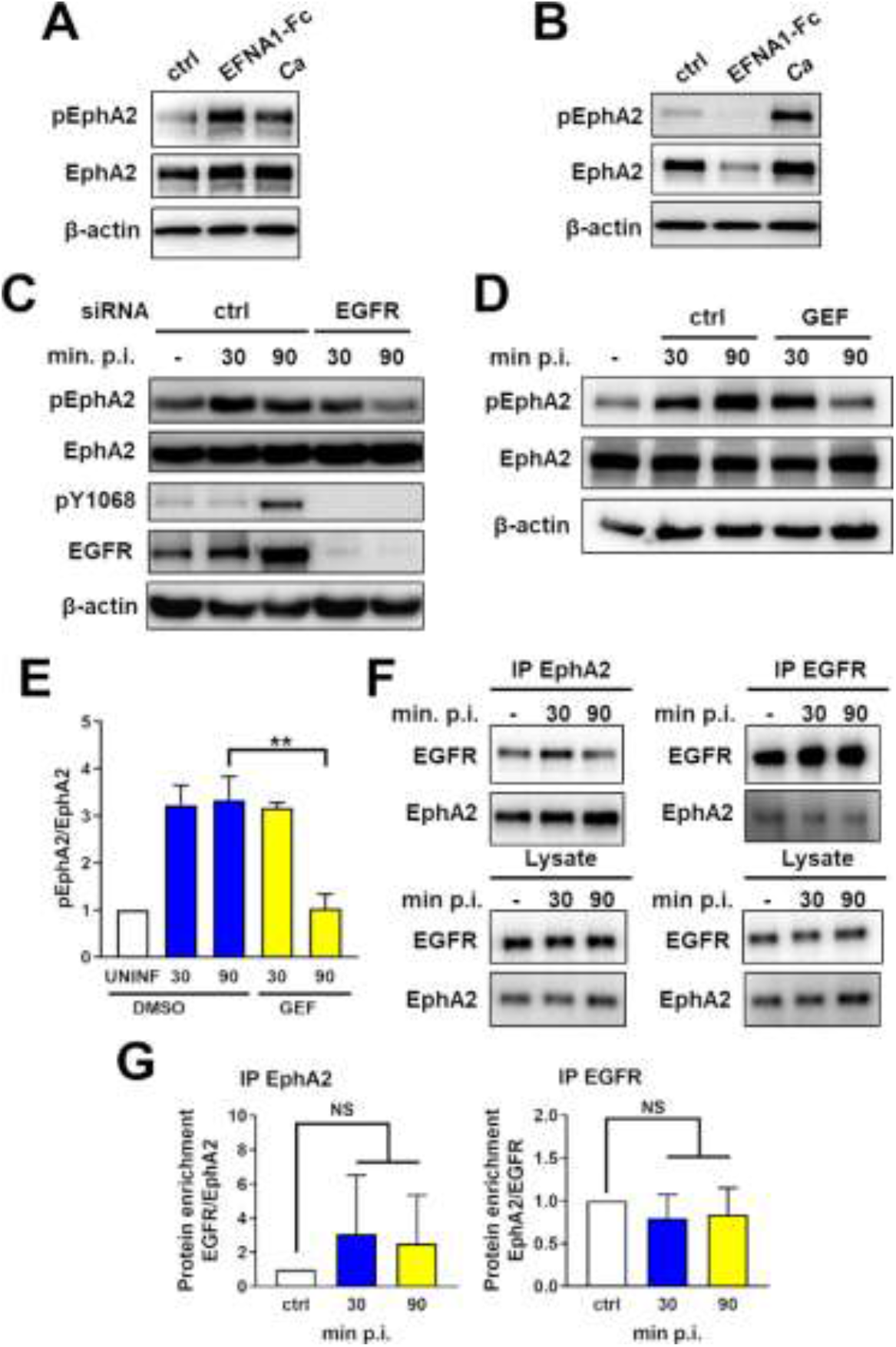
EGFR activity is required for sustained EphA2 phosphorylation. (A and B) Immunoblots showing effects of ephrin A1-Fc (EFNA1-Fc) or yeast-phase *C. albicans* SC5314 (Ca) on the phosphorylation of ephrin type-A receptor 2 (EphA2) in OKF6/TERT-2 oral epithelial cells after stimulation for 15 min (A) and 60 min (B) post-infection (p. i.). Results are representative of 3 independent experiments. Densitometric quantification of all 3 immunoblots is shown in S1 Fig. (C and D) Effects of epidermal growth factor (EGFR) siRNA (C) and the EGFR kinase inhibitor gefitinib (GEF) (D) on the time course of EphA2 and EGFR phosphorylation in oral epithelial cells infected with *C. albicans*. Results are representative of 3 independent experiments. Densitometric quantification of all 3 immunoblots such as the one in Fig. 1C is shown in S1 Fig. (E) Densitometric quantification of all 3 immunoblots such as the one in Fig. 3D. (F) Lysates of oral epithelial cells infected with *C. albicans* for 30 and 90 min were immunoprecipitated (IP) with antibodies against EphA2 (left) and EGFR (right), after which EphA2 and EGFR were detected by immunoblotting (Top). Immunoblots of lysates prior to immunoprecipitation, demonstrating equal amounts of input protein (Bottom). (G) Densitometric analysis of 3 independent immunoblots such as the ones shown in (F). Results are mean ± SD. Statistical significance relative to uninfected control cells was analyzed by the Student’s t test assuming unequal variances. **, *p* < 0.01; NS, not significant.

### EGFR activation sustains *C. albicans-induced* EphA2 phosphorylation

EphA2 and EGFR function in the same pathway to mediate the endocytosis of *C. albicans* by oral epithelial cells *in vitro*. Also, EphA2 is required for *C. albicans* to activate EGFR because siRNA knockdown of EphA2 in oral epithelial cells blocks fungal-induced phosphorylation of EGFR [16]. Cross-talk between EphA2 and EGFR has also been observed in other cell types [20, 21]. To determine if EGFR is required for sustained activation of EphA2, we tested the effects of inhibiting EGFR on *C. albicans*-induced EphA2 phosphorylation. When EGFR was either knocked down with siRNA or inhibited with the specific EGFR kinase inhibitor, gefitinib [22], EphA2 phosphorylation was transient, occurring within 30 min of infection, but declining to basal levels by 90 min (Fig 1C-1E, S1D-S1F Fig). Therefore, activation of EGFR by *C. albicans* is required to sustain EphA2 phosphorylation.

Cross talk between EphA2 and EGFR may reflect a physical interaction between these two receptors. We investigated this possibility by immunoprecipitation experiments. When EphA2 was immunoprecipitated from lysates of uninfected epithelial cells, EGFR was pulled down with it (Fig 1F and 1G). The interaction between EphA2 and EGFR appeared to be constitutive because the amount of EGFR that was associated with EphA2 did not change significantly when the epithelial cells were infected with *C. albicans*. In reciprocal experiments, we determined that immunoprecipitation of EGFR from infected and uninfected epithelial cells also pulled down similar amounts of EphA2 (Fig 1F and 1G). These results indicate that EphA2 and EGFR likely form part of a complex that enables each receptor to influence the activity of the other.

### *C. albicans* Als3 and Ece1 are required for prolonged EphA2 activation

Two *C. albicans* virulence factors, the Als3 invasin and the Ece1/Candidalysin toxin, have been shown to be required for activation of EGFR [18, 23, 24]. To investigate the relationship between Als3 and Ece1/Candidalysin and the stimulation of epithelial cells, we constructed *als3Δ/Δ* and *ece1*Δ/Δ single gene deletion mutants and an *als3*Δ/Δ *ece1*Δ/Δ double gene deletion mutant. As expected [3, 7], when epithelial cells were infected with either the *als3*Δ/Δ or *als3*Δ/Δ *ece1*Δ/Δ mutants, the number of cell-associated *C. albicans* cells (a measure of adherence) and endocytosed *C. albicans* cells was significantly reduced relative to the wild-type strain (Fig 2A). By contrast, the *ece1*Δ/Δ mutant acted similarly to the wild-type parent strain in these assays. Also, both the *als3*Δ/Δ and *ece1*Δ/Δ mutants caused significantly less epithelial cell damage than the wild-type strain (Fig 2B). The *als3Δ/Δ ece1Δ/Δ* double mutant caused the least amount of epithelial cell damage, which was significantly less than that caused by the *als3*Δ/Δ single mutant (Fig 2B). Collectively, these data support the model that Als3-mediated epithelial cell invasion is required for Ece1-induced damage.

**Fig 2.**
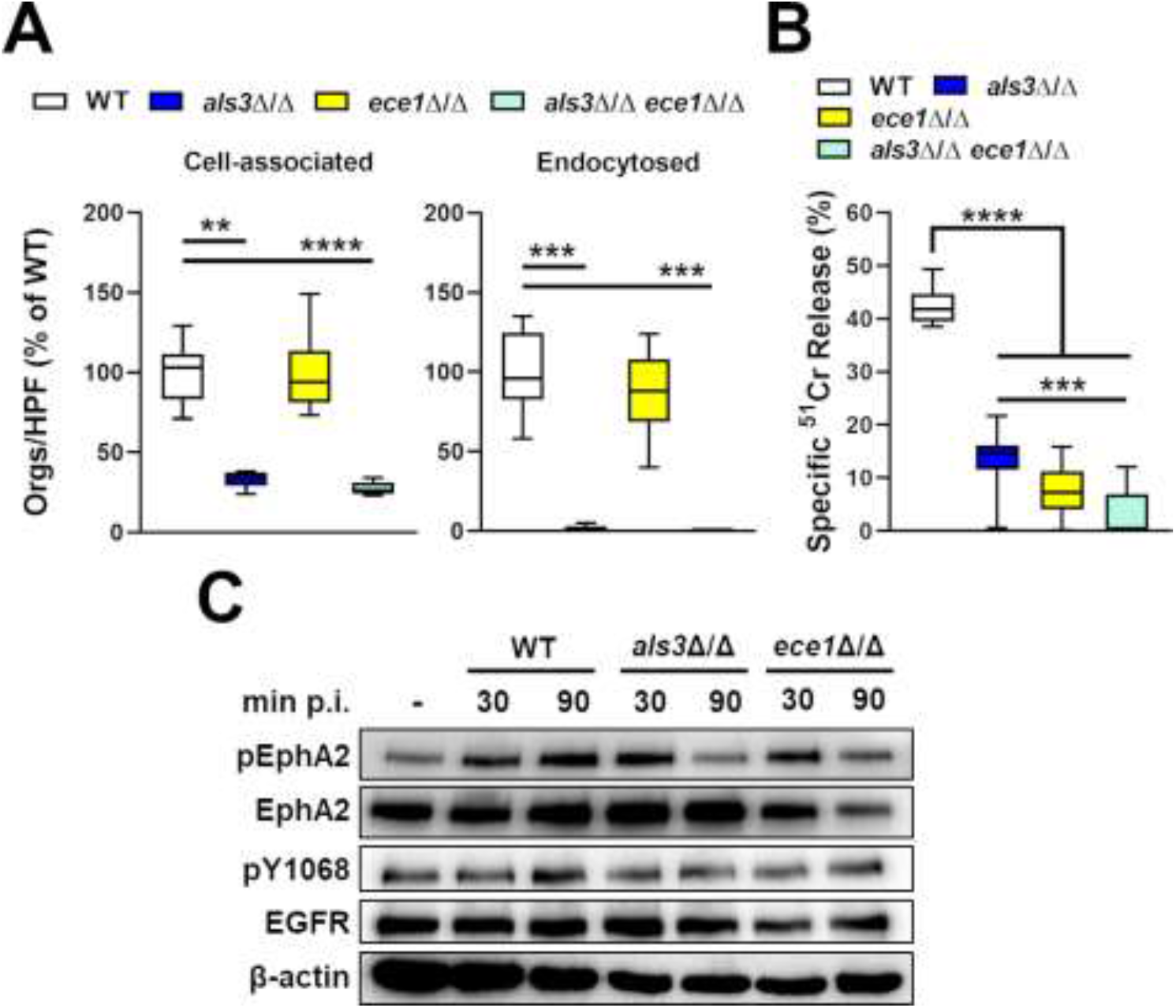
Effects of *C. albicans als3*Δ/Δ and *ece1*Δ/Δ mutants EGFR and EphA2 activation. (A) The number of cell-associated (a measure of adherence) and endocytosed organisms (orgs) per high-power field (HPF) of the indicated *C. albicans* strains after 120 min of infection of OKF6/TERT-2 oral epithelial cells. (B) Extent of epithelial cell damage caused by the indicated strains after 8 h of infection. Data in (A and B) are the combined results of three experiments, each performed in triplicate. Statistical significance was determined by analysis of variance with Dunnett’s test for multiple comparisons. **, *p* < 0.01; ***, *p* < 0.001; ****, *p* < 0.0001. (C) Immunoblot analysis showing the effects of the indicated *C. albicans* strains on the phosphorylation of EphA2 and EGFR after 30 and 90 min post-infection (p. i.). Result are representative a 3 independent experiments. Densitometric quantification of all 3 immunoblots such as the one in Fig. 3C is shown in S2 Fig.

To determine if Als3 and Ece1 are required for sustained EphA2 activation, we analyzed the phosphorylation of EphA2 and EGFR in response to the *als3*Δ/Δ and *ece1*Δ/Δ mutant strains. While infection with both mutant strains induced EphA2 phosphorylation in oral epithelial cells after 30 min, neither mutant strain was able to sustain EphA2 phosphorylation at 90 min (Fig 2C, S2 Fig). This lack of sustained EphA2 phosphorylation was likely due to the absence of EGFR activation, because neither the *als3*Δ/Δ nor the *ece1*Δ/Δ mutant induced significant phosphorylation of EGFR.

### EGFR induces a subset of EphA2-mediated pro-inflammatory responses in oral epithelial cells

To investigate the roles of EGFR and EphA2 in stimulating an epithelial cell proinflammatory response *in vitro*, we used siRNA to knock down EGFR and EphA2 prior to infecting the cells with *C. albicans*. We found that knockdown of EGFR inhibited the secretion of CXCL8/IL-8 and CCL20 similarly to knockdown of EphA2 (Fig 3A). Although knockdown of EGFR had no effect on the release of IL-1α, and IL-1β, knockdown of EphA2 reduced the secretion of both of these cytokines. Inhibition of EGFR phosphorylation with gefitinib also inhibited the secretion of CXCL8/IL-8 and CCL20, but not IL-1α or IL-1β (Fig 3B). These results suggest that activation of both EphA2 and EGFR is required for *C. albicans* to induce epithelial cells to secrete CXCL8/IL-8 and CCL20. They also indicate that EphA2 mediates production of IL-1α and IL-1β independently of EGFR.

**Fig 3.**
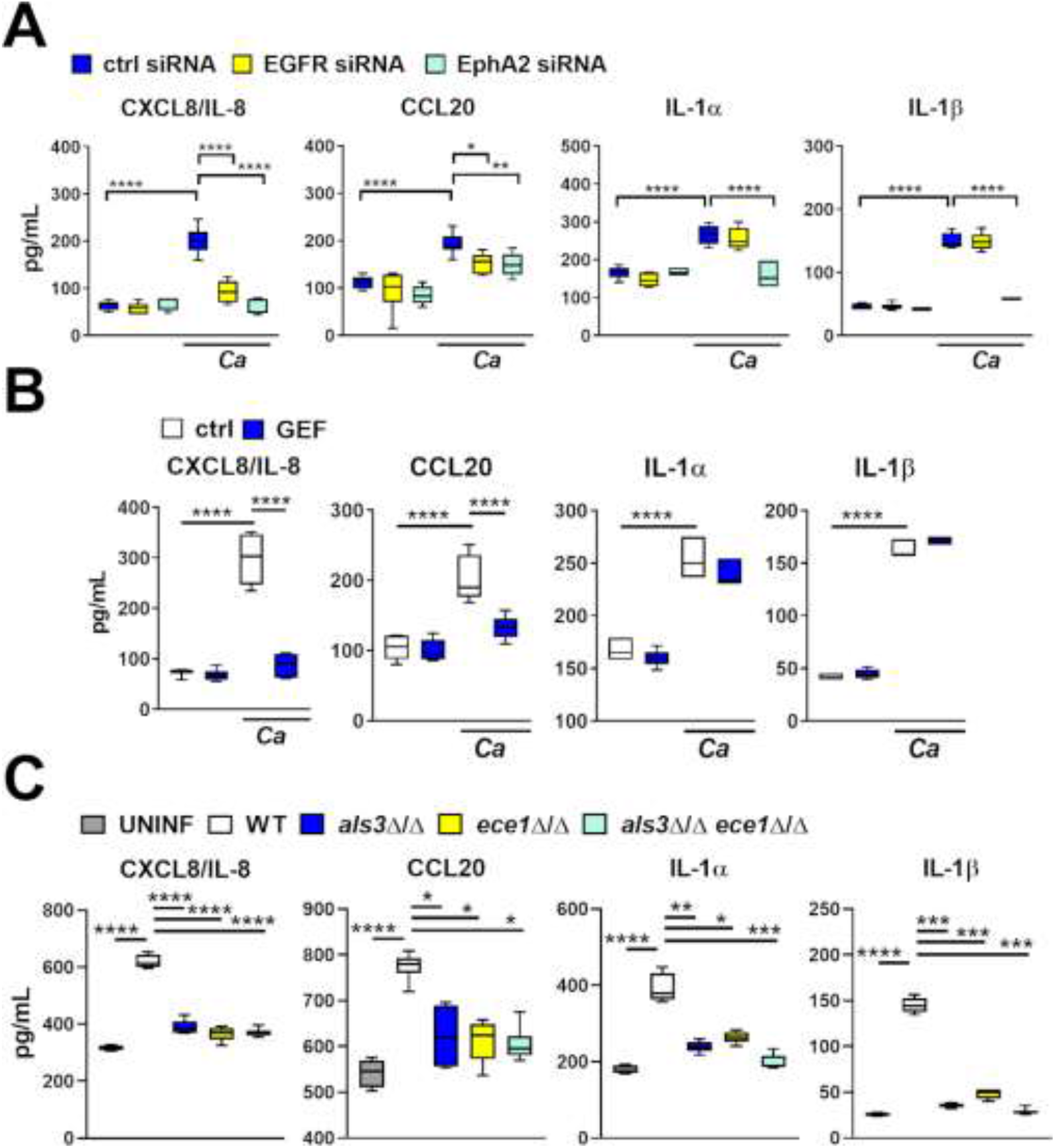
EGFR signaling is required to induce a subset of epithelial cells inflammatory responses *in vitro*. (A) Effects of siRNA knockdown of EGFR and EphA2 on the production of the indicated cytokines and chemokines by OKF6/TERT-2 oral epithelial cell in response to 8 h of infection with *C. albicans*. (B) Effects of the EGFR inhibitor gefitinib on the epithelial cell response to *C. albicans*. (C) Stimulation of epithelial cells by the indicated strains of *C. albicans*. Box whisker plots show median, interquartile range, and range of 3 independent experiments, each performed in duplicate. The data were analyzed using the Kruskal-Wallis corrected for multiple comparisons. *, *p* < 0.05; **, *p* < 0.01; ***, *p* < 0.001; \*\*\*\**p*, < 0.0001; *Ca*, *C. albicans;* ctrl, control; GEF, gefitinib.

Next, we investigated the roles of Als3 and Ece1 in stimulating oral epithelial cells *in vitro*. Infection with the *als3*Δ/Δ and *ece1*Δ/Δ single mutants and the *als3*Δ/Δ *ece1*Δ/Δ double mutant induced significantly less secretion of CXCL8/IL-8, CCL20, IL-1α, and IL-1β than the wild-type parent strain (Fig 3C). Of note, the epithelial cell stimulation defect of the *als3*Δ/Δ *ece1*Δ/Δ double mutant was no greater than that of either the *als3*Δ/Δ or *ece1*Δ/Δ single mutant. Collectively, these results support the model that *C. albicans* Als3 and Ece1/Candidalysin function in the same pathway to activate EGFR, which prolongs EphA2 phosphorylation and stimulates oral epithelial cells to secrete CXCL8/IL-8 and CCL20. These results also suggest that Als3 and Ece1/Candidalysin induce the production of IL-1α and IL-1β via a mechanism that requires EphA2 but is independent of EGFR.

### The response of OKF6/TERT-2 epithelial cells to *C. albicans* differs significantly from TR146 epithelial cells

The above experiments were conducted with OKF6/TERT-2 oral epithelial cells, which were generated by transfecting normal human cells with a constitutively active telomerase [25]. Using the TR146 human buccal squamous cell carcinoma cell line, Ho et al. also found that *C. albicans*-induced activation of EGFR is required for the fungus to stimulate oral epithelial cells to secrete multiple cytokines [18]. One difference between their results and those presented here is that EGFR was required for induction of IL-1α and IL-1β in TR146 cells but not OKF6/TERT-2 cells. To investigate this difference, we compared the interactions of *C. albicans* with both cell lines. We found that while the adherence of *C. albicans* to both types of epithelial cells was similar, TR146 cells endocytosed over 3-fold more *C. albicans* cells than did OKF6/TERT-2 cells (Fig 4A). Also, gefitinib modestly inhibited the adherence of *C. albicans* to TR146 cells, but not OKF6/TERT-2 cells, and it inhibited *C. albicans* endocytosis by TR146 cells by 30% and endocytosis by OKF6/TERT-2 cells by 64%. Thus, in TR146 cells relative to OKF6/TERT-2 cells, EGFR is more important for adherence but less important for inducing the endocytosis of *C. albicans*.

**Fig 4.**
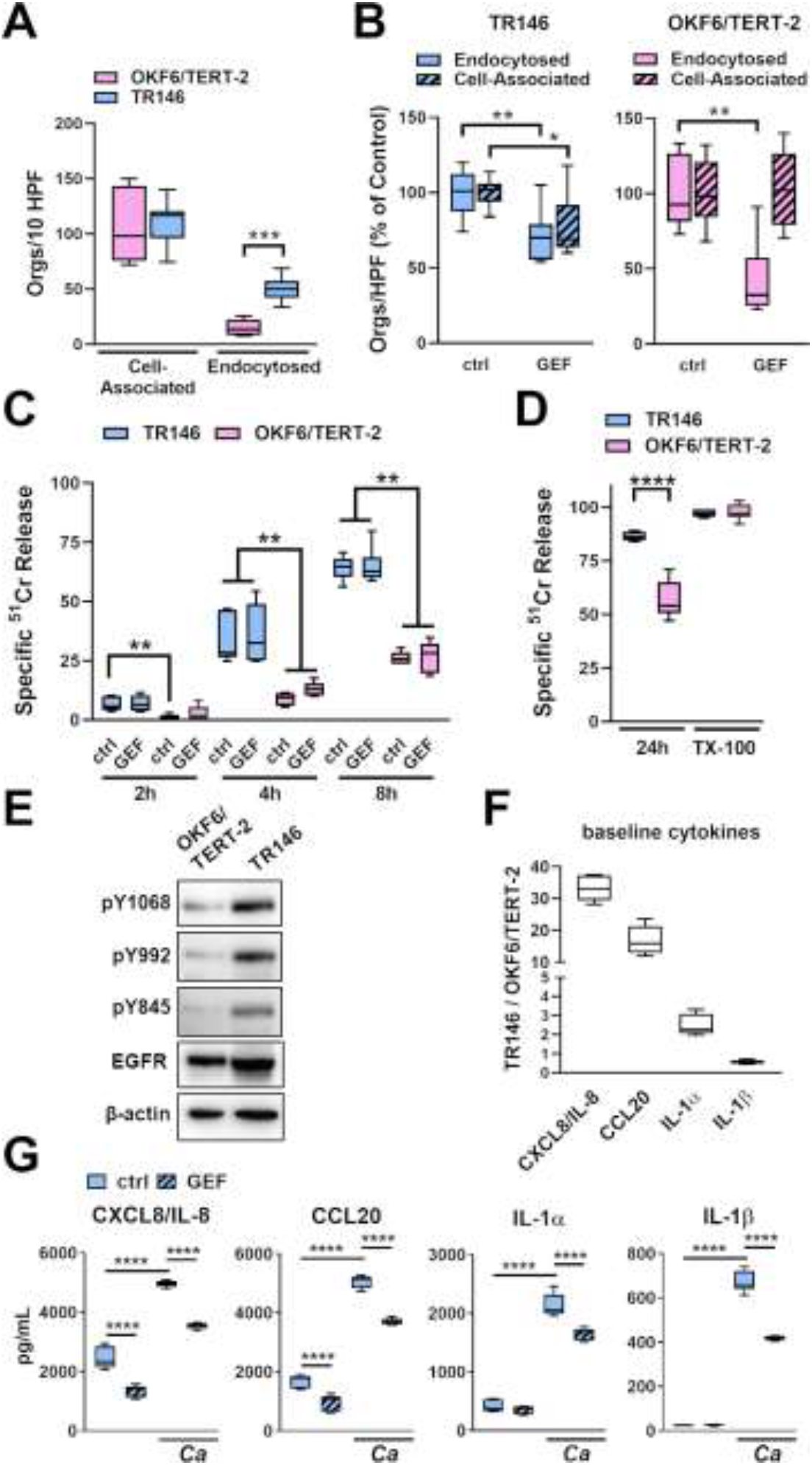
Comparison of the response of two oral epithelial cell lines to *C. albicans* infection and EGFR inhibition. (A) Adherence and endocytosis of wild-type *C. albicans* by OKF6/TERT-2 and TR146 oral epithelial cell lines. (B) Comparison of the effects of gefitinib (GEF) on *C. albicans* adherence to and endocytosis by OKF6-TERT-2 and TR146 cells. (C) Time course of *C. albicans*-induced damage to OKF6-TERT-2 and TR146 cells. (D) Extent of damage to OKF6-TERT-2 and TR146 cells caused by *C. albicans* and Triton X-100 after 24 h incubation. (E) Comparison of the extent of phosphorylation on the indicated tyrosine residues of EGFR phosphorylation in unstimulated OKF6/TERT-2 and TR146 cells. Shown in a representative immunoblot from 3 independent experiments. (F) Comparison of the basal release of the indicated inflammatory mediators by uninfected OKF6-TERT-2 and TR146 cells. Results are the ratio of TR146 cells to OKF6/TERT2 cells. (G) Effects of gefitinib on *C. albicans-induced* production of the indicated pro-inflammatory mediators by TR146 cells. Graphs show the results of 3 experiments, each performed in triplicate (A-D) or duplicate (F and G). The data in (A, B, and D) were analyzed with the Mann-Whitney test, and the data in (C and G) were analyzed by the Kruskal-Wallis test corrected for multiple comparisons. *, *p* < 0.05; **, *p* < 0.01; ***, *p* < 0.001; \*\*\*\**p*, < 0.0001; *Ca*, *C. albicans;* ctrl, control; GEF, gefitinib; TX-100, Triton X-100.

We also used a ^51^Cr release assay to compare the susceptibility of the two cell types to damage induced by *C. albicans*. In a time-course study, *C. albicans* caused significantly more damage to TR146 cells than to OKF6/TERT-2 cells at all time points (Fig 4C). In fact, after 24 hr of infection, *C. albicans-induced* damage of TR146 cells was near-maximal, whereas the extent of damage to the OKF6/TERT-2 cells was significantly less (Fig 4D). Gefitinib had no effect on the extent of *C. albicans-induced* damage to either cell line. Collectively, these data indicate that TR146 cells are more susceptible to *C. albicans-induced* damage than OKF6/TERT-2 cells.

The extent of EGFR phosphorylation and cytokine production by TR146 cells and OKF6/TERT-2 cells also differed. Uninfected TR146 cells had significantly higher levels of total and phosphorylated EGFR than OKF6/TERT-2 cells (Fig 4E, S3 Fig). Uninfected TR146 cells also secreted over 30-fold more CXCL8/IL-8, 15-fold more CCL20, and 2-fold more IL-1α than OKF6/TERT-2 cells (Fig 4F). The levels of these cytokines and IL-1β significantly increased when the TR146 cells were infected with *C. albicans* (Fig 4G). While gefitinib inhibited the production of CXCL8/IL-8 and CCL20 by *C. albicans* infected TR146 cells, it also decreased the production of IL-1α and IL-1β (Fig 4G). Thus, EGFR governs *C. albicans-induced* production of chemokines in both cell lines, but it regulates production of IL-1α and IL-1β only in TR146 cells.

### Inhibition of EGFR decreases EphA2 phosphorylation, oral fungal burden, and the host inflammatory response during OPC

To assess the role of EGFR in mediating the host inflammatory response to *C. albicans* during OPC, we treated immunocompetent mice with gefitinib and then orally inoculated them with *C. albicans*. Immunocompetent mice were used to avoid potentially confounding effects of immunosuppression on the host inflammatory response. When immunocompetent mice are orally inoculated with *C. albicans* strain SC5314, infection persists for 2 days, after which the organism is cleared by the host inflammatory response [14, 26].

Using flow cytometry and phosphospecific antibodies, we assessed the effects of gefitinib on EGFR and EphA2 phosphorylation in the oral epithelium. As predicted by the *in vitro* data, treatment with gefitinib reduced the phosphorylation of both EGFR and EphA2 in CD45^-^ EpCam^+^ cells after 1 d of infection (Fig 5). At this time point, the gefitinib-treated mice had a 3-fold reduction in oral fungal burden compared to untreated mice (Fig 6A), probably due to inhibition of epithelial cell invasion by the drug.

**Fig 5.**
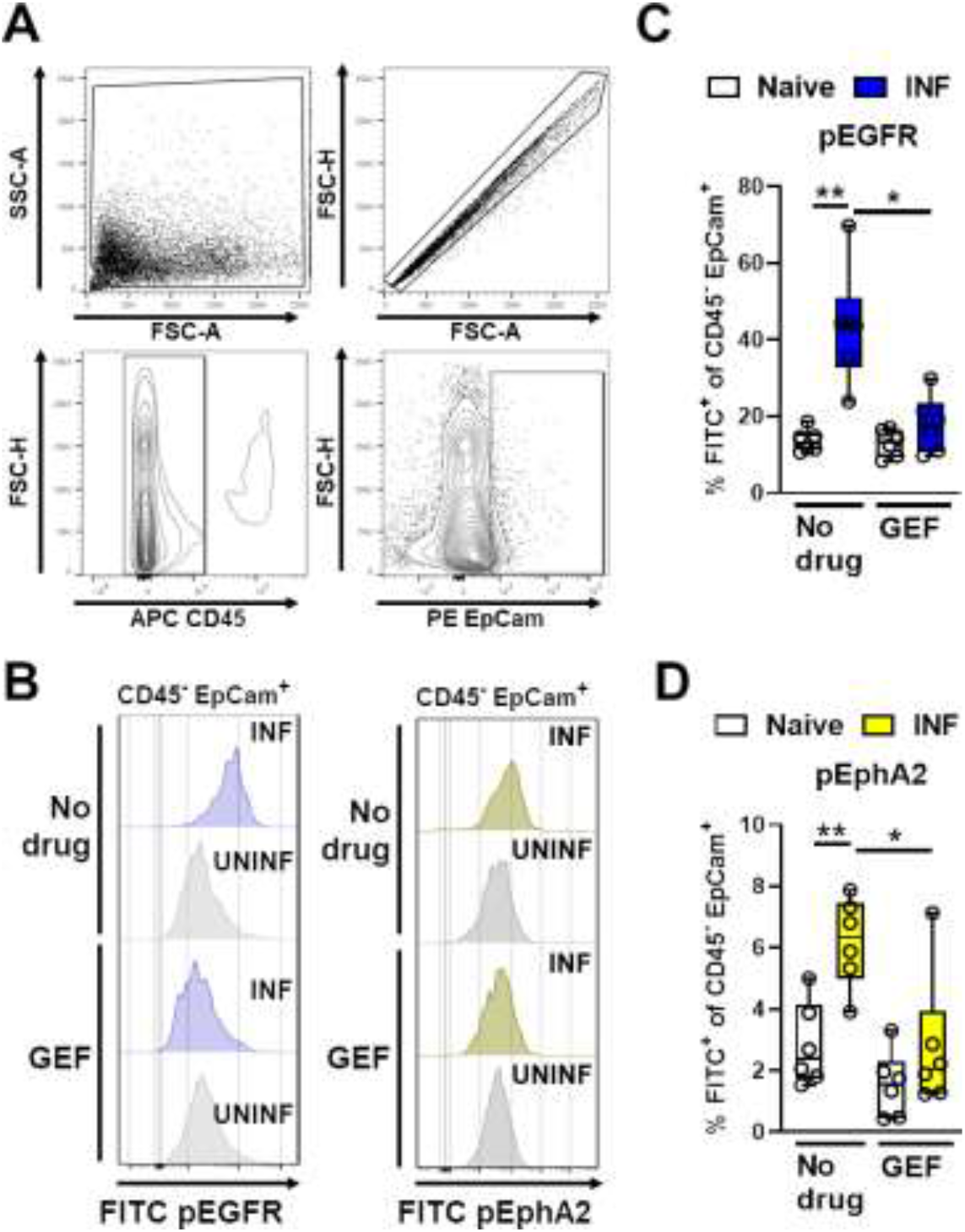
Pharmacological inhibition of EGFR reduces *C. albicans-induced* EphA2 activation during OPC. (A) Gating strategy to determine EGFR and EphA2 phosphorylation in the oral epithelial cells of mice with OPC. (B) Representative histograms of CD45^-^ EpCam^+^ cells showing the effects of *C. albicans* infection and gefitinib (GEF) treatment on the phosphorylation of EGFR and EphA2 after 1 d of OPC. (C-D) Effects of gefitinib on the percentage of oral epithelial cells with phosphorylated EGFR (C) and EphA2 (D) in mice after 1 d of OPC. Data are combined results from 6 mice per group from a single experiment. Statistical significance was determined using the Mann-Whitney test. *, *p* < 0.05; **, *p* < 0.01; INF, infected; UNINF, uninfected.

**Fig 6.**
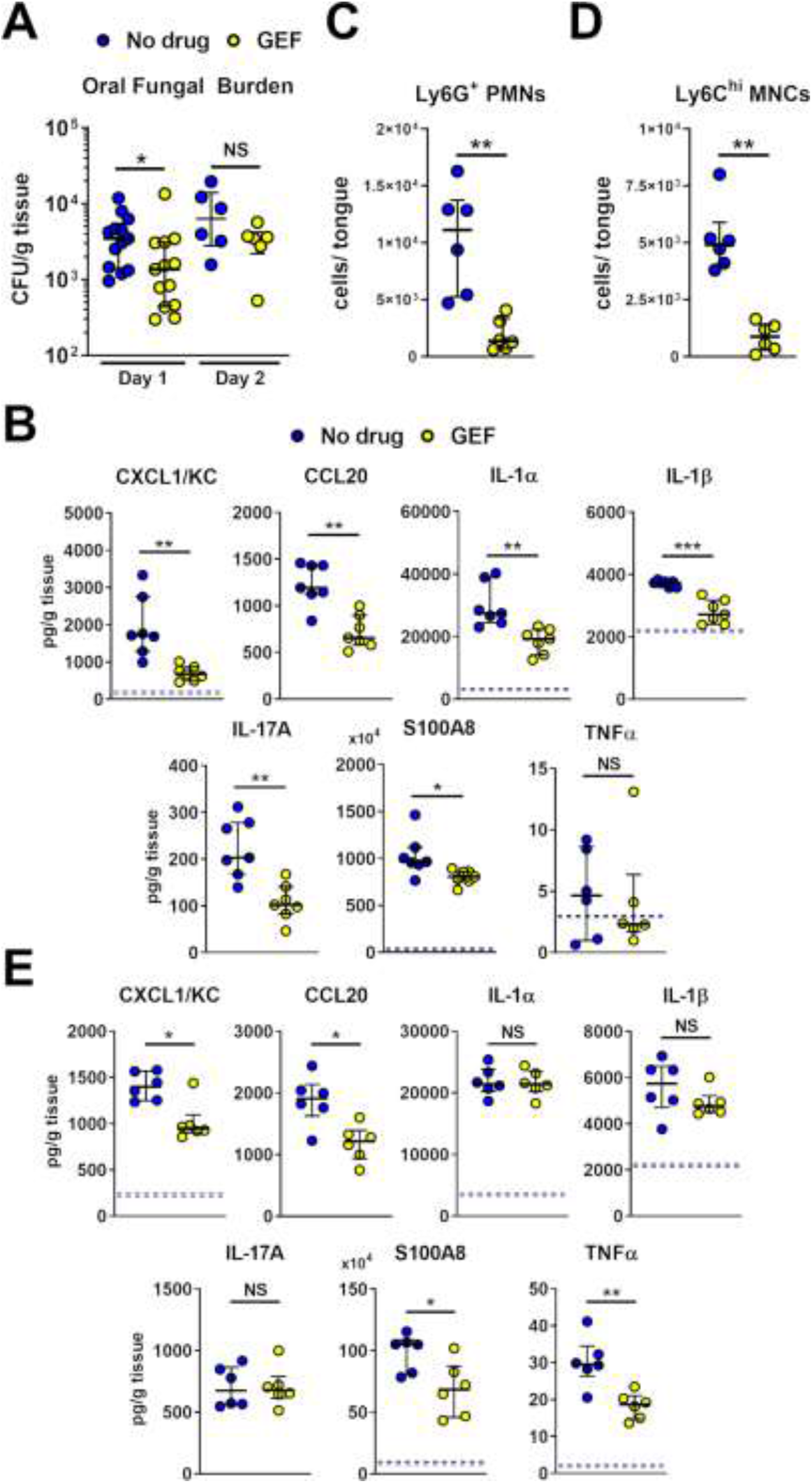
Pharmacological inhibition of EGFR decreases early fungal burden and reduces the inflammatory response. (A) Oral fungal burden of control (no drug) and gefitinib (GEF) treated mice after 1 and 2 d post-infection. Results are the median ± interquartile range of at total of 14 mice per group from two independent experiments on d 1 and of a total of 6 mice per group from a single experiment on day 2. The y-axis is set at the limit of detection (20 CFU/g tissue). (B and E) Levels of the indicated inflammatory mediators in the oral tissues of mice after 1 d (B) and 2 d(E) of infection. Results are from at total of 7 mice per group from two independent experiments on d 1 and from a total of 6 mice per group from a single experiment on d 2. Dashed line indicates the median level of inflammatory mediators in uninfected mice if above 0. (C and D) Levels of neutrophils (C) and inflammatory monocytes (D) in the tongues of control and gefitinib-treated mice after 1 d of infection. Results are from a total of 6 mice per group from two independent experiments. Statistical significance in (A, C and D) was determined using the Mann-Whitney test and in (B and E) by the Kruskal-Wallis test corrected for multiple comparisons. NS, not significant; *, *p* < 0.05; **, *p* < 0.01.

To determine the effects of gefitinib on the inflammatory response, we measured whole tongue levels of CXCL1/KC, CCL20, IL-1α and IL-1β. To obtain a more comprehensive view of the effects of gefitinib, we also measured the levels of IL-17A, S100A8, and TNFα. After 1 day of infection, the oral tissues of the gefitinib-treated mice contained significantly less CXCL1/KC, CCL20, IL-1α, IL-1β, IL-17A, and the host defense peptide S100A8 than control mice (Fig 6B). There were also significantly fewer neutrophils and inflammatory monocytes in the oral tissues of the gefitinib-treated mice (Fig 6C and 6D, S4 Fig). After 2 days of infection, the gefitinib-treated and control mice had similar oral fungal burden (Fig 6A). At this time point, treatment with gefitinib reduced the levels of CXCL1/KC, CCL20, S100A8 and TNFα, but not IL-1α, IL-1β, and IL-17A (Fig 6E). These results indicate that blocking EGFR reduces a subset of the early inflammatory response to acute OPC in immunocompetent mice.

Because gefitinib impaired the host inflammatory response to *C. albicans*, we investigated its effects on the candidacidal activity of neutrophils and macrophages. We found that gefitinib did not decrease the capacity of human neutrophils, bone marrow (BM) neutrophils isolated from gefitinib-treated mice, or murine BM derived macrophages to kill *C. albicans in vitro* (S5 Fig). Taken together, these results indicate that while EGFR signaling is required for epithelial cells to mount a pro-inflammatory response to *C. albicans*, it is dispensable for governing phagocyte killing of the organism.

### *C. albicans* adhesins/invasins and Ece1/Candidalysin function together during OPC

To investigate the roles Als3 and Ece1/Candidalysin in stimulating the host response during oropharyngeal candidiasis, we orally infected immunocompetent mice with *C. albicans als3Δ/Δ* and *ece1*Δ/Δ mutants. After 1 day of infection, mice inoculated with the *als3*Δ/Δ mutant had reduced oral fungal burden relative to those infected with the wild-type strain, and mice inoculated with the *ece1*Δ/Δ mutant had an even greater decrease in fungal burden (Fig 7A). After 2 days of infection, the fungal burden of mice infected with the *als3*Δ/Δ mutant was similar to that of mice infected with the wild-type strain, whereas the fungal burden of mice infected with the *ece1*Δ/Δ mutant remained significantly lower (Fig 7A). We also determined that C57BL/6 mice infected with the *ece1*Δ/Δ mutant constructed in either strain SC5314 or BWP17 had reduced oral fungal burden (S6 Fig). Thus, in the acute oropharyngeal candidiasis model, Als3 is necessary for maximal infection at day 1, whereas Ece1/Candidalysin is required for maximal infection at both days 1 and 2.

**Fig 7.**
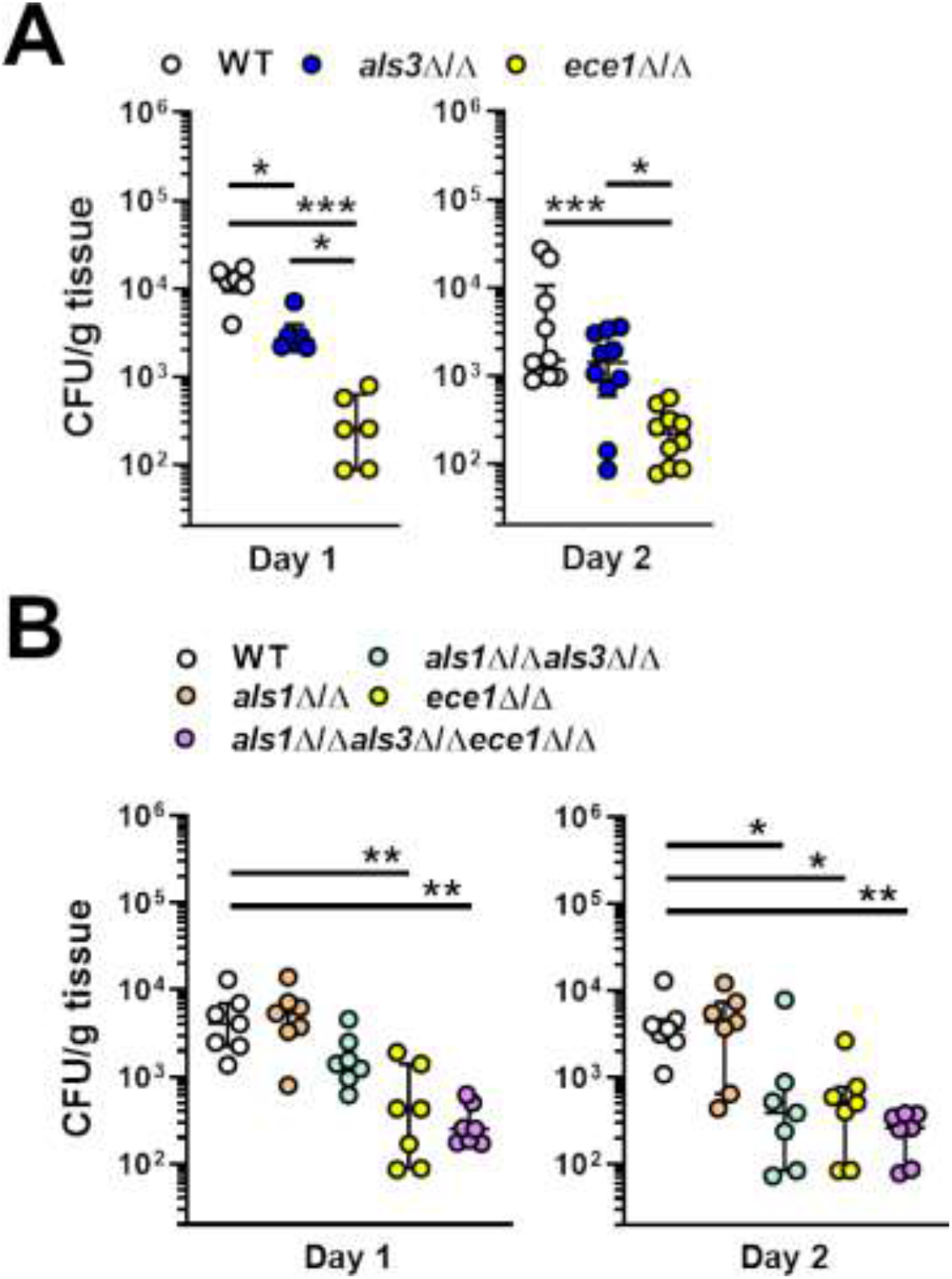
Roles of Als1, Als3, Ece1/Candidalysin in *C. albicans* virulence during OPC. (A and B) Oral fungal burden of Balb/c mice 1 and 2 d after inoculation with the indicated strains of *C. albicans*. Results are the median ± interquartile range of 6-8 mice per strain from single experiments, except for d 2 of in (A), which are from a total of 10 mice from 2 independent experiments. The data were analyzed by the Kruskal-Wallis test corrected for multiple comparisons. *, *p* < 0.05; **, *p* < 0.01; ***, *p* < 0.001.

*C. albicans* must first adhere to host cells in order to invade them. *C. albicans* Als1 is structurally similar to Als3 and it can function as both an adhesin and invasin [3, 27]. We reasoned that the presence of Als1 in *als3*Δ/Δ mutant could enable the organism to adhere to and invade the oral epithelium in the absence of Als3. To test this hypothesis, we constructed *als1*Δ/Δ, *als1*Δ/Δ *als3*Δ/Δ, and *als1*Δ/Δ *als3*Δ/Δ *ece1*Δ/Δ mutants. *In vitro*, the *als1*Δ/Δ *als3*Δ/Δ mutant had significantly reduced adherence to OKFG/TERT-2 cells compared to the *als3*Δ/Δ mutant, indicating that Als1 and Als3 make independent contributions to *C. albicans* adherence (S7A Fig). Because the *als3*Δ/Δ single mutant was endocytosed so poorly by oral epithelial cells and caused so little host cell damage, deletion of *ALS1* did not reduce these interactions further (S8B and S8C Fig)

After 1 day of oral infection, mice infected with the *als1Δ/Δ als3Δ/Δ* mutant had a slight reduction in oral fungal burden relative to mice infected with the wild-type strain, but this difference was not statistically significant (Fig 7B). At this time point, the fungal burden of mice infected with *ece1*Δ/Δ single mutant and the *als1Δ/Δ als3Δ/Δ ece1Δ/Δ* triple mutant was significantly reduced. After 2 days of infection, the fungal burden of mice infected with the *als1Δ/Δ als3Δ/Δ* double mutant was decreased to the same extent as mice infected with either the *ece1*Δ/Δ single mutant or the *als1*Δ/Δ *als3*Δ/Δ *ece1*Δ/Δ triple mutant (Fig 8B). These data suggest that adherence and invasion mediated by Als1 and Als3 combined with host cell damage caused by Ece1/Candidalysin play key roles in inducing acute OPC.

**Fig 8.**
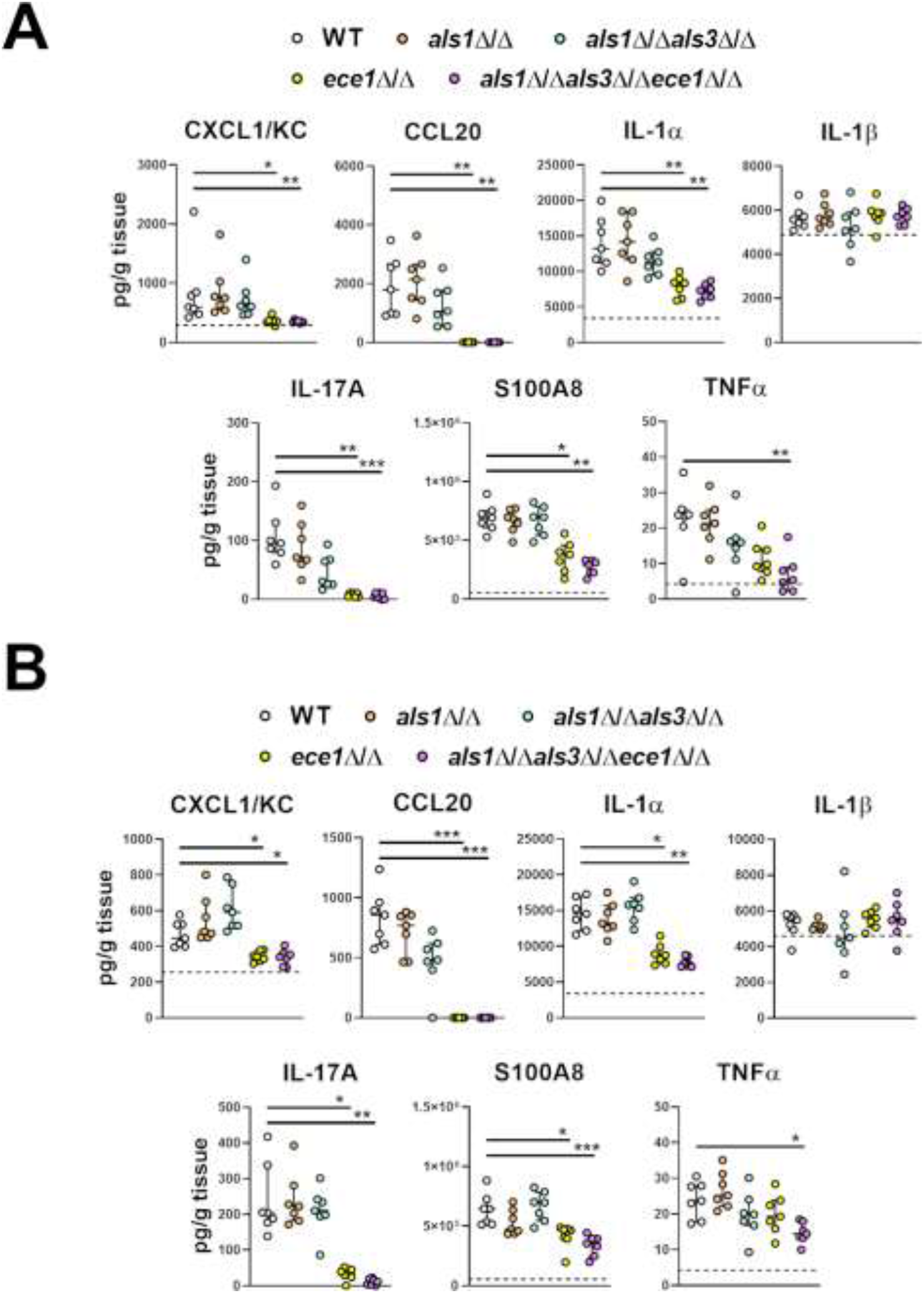
Induction of the oral inflammatory response by *C. albicans* invasins and Ece1/Candidalysin. Levels of the indicated inflammatory mediators in the tongues of mice orally inoculated with indicated strains of *C. albicans* after 1 d (A) and 2 d (B) of infection. Results are the median ± interquartile range of a total of 7 mice in each group from a single experiment. Median levels of inflammatory mediators in uninfected mice are indicated with dashed line if above 0. The data were analyzed by the Kruskal-Wallis test corrected for multiple comparisons. *, *p* < 0.05; **, *p* < 0.01; ***, *p* < 0.001.

To determine the contributions of Als1, Als3 and Ece1/Candidalysin to the induction and maintenance of the oral innate immune response, we determined the levels of inflammatory mediators in the infected tongues after 1 and 2 days of infection. Three different patterns of response were observed. The levels of IL-1β were similar in mice infected with all strains of *C. albicans*, indicating that the production of this cytokine is independent of Als1, Als3 and Ece1/Candidalysin (Fig 8). The levels of many inflammatory mediators, such as IL-1α, CXCL1/KC CCL20, IL-17A, and S100A8 were decreased to the same extent in mice infected with either the *ece1*Δ/Δ mutant or the *als1*Δ/Δ *als3*Δ/Δ *ece1*Δ/Δ mutant, suggesting that the production of these mediators is dependent on Ece1/Candidalysin but independent of Als1 and Als3. The levels of TNFα were significantly reduced only in mice infected with the *als1*Δ/Δ *als3*Δ/Δ *ece1*Δ/Δ mutant, signifying that Als1, Als3, and Ece1/Candidalysin function cooperatively to induce the production of this cytokine. Thus, while the production of some inflammatory mediators during OPC is largely driven by Ece1/Candidalysin, Als1 and Als3 contribute to the production of TNFα.

## Discussion

Oral epithelial cells play a central role in orchestrating the host defense against *C. albicans* during OPC [1, 11, 28, 29]. Our current data show that in oral epithelial cells, EphA2 and EGFR form part of a complex and that EGFR activation mediates a subset of proinflammatory response to this fungus. Previously, we found that EphA2 activation is required for *C. albicans* to stimulate EGFR [16]. Here, we determined that activation of EGFR is in turn necessary for *C. albicans* to sustain activation of EphA2, both *in vitro* and *in vivo*. Thus, these two receptors function interdependently to mediate the epithelial cell response to *C. albicans*. These data also indicate that the initiation of a proinflammatory response in oral epithelial cells requires two signals, one induced by EphA2 binding to fungal β-glucans and the other induced by EGFR stimulated by fungal invasins and Ece1/Candidalysin.

EphA2 and EGFR are known to interact in epithelial cell cancers, especially those that have become resistant to EGFR inhibitors, in which siRNA knockdown of EphA2 restores sensitivity to pharmacological EGFR inhibition [30, 31]. Interestingly, treatment of malignant cell lines with soluble EFNA1 has the same effect as EphA2 siRNA, presumably because EFNA1 induces EphA2 endocytosis and subsequent degradation. Our data indicate that *C. albicans* activates EphA2 differently than EFNA1 because binding of *C. albicans* stabilizes EphA2 and prevents its degradation. The sustained EphA2 protein levels likely contribute to the prolonged EphA2 signaling induced by *C. albicans* infection.

We found that *C. albicans* activation of EGFR resulted in the production of proinflammatory mediators by two different oral epithelial cell lines. Recently, it has been reported that Ece1/Candidalysin activates EGFR and stimulates the TR146 oral epithelial cell line to produce Il-1α, IL-1β, IL-6, G-CSF, and GM-CSF [18]. In the current work, knockdown of EGFR with siRNA or inhibition of EGFR with gefitinib significantly reduced *C. albicans-induced* production of CXCL8/IL-8 and CCL20 by both TR146 cells and OKF6-TERT2 cells. However, inhibition of EGFR only decreased production of Il-1α and IL-1β by TR146 cells, but not by OKF6-TERT2 cells. It was also determined that TR146 cells had higher basal levels of total and phosphorylated EGFR than OKF6-TERT2 cells, which may explain why cytokine production by TR146 cells was more sensitive to EGFR inhibition. Nevertheless, the result with both oral epithelial cells lines indicate that *C. albicans* activates EGFR, which induces the production of proinflammatory mediators.

The current data indicate a functional interaction between Als3 and Ece1/Candidalysin. Consistent with previous reports [3, 7], we found that both the *als3*Δ/Δ mutant and the *ece1*Δ/Δ mutant were defective in damaging oral epithelial cells *in vitro*. Also, these two mutants were similarly impaired in their capacity to stimulate epithelial cells to secrete proinflammatory mediators. Strikingly, deletion of *ALS3* in the *ece1*Δ/Δ mutant did not lead to a further reduction in epithelial cell damage and production of proinflammatory mediators. These results indicate that Als3 and Ece1/Candidalysin function in the same pathway to damage epithelial cells and stimulate them to secrete proinflammatory mediators *in vitro*. A potential explanation for the functional linkage between Als3 and Ece1/Candidalysin is that when *C. albicans* hyphae are endocytosed via Als3, secreted Ece1/Candidalysin is contained within the endocytic vacuole, where it accumulates to sufficiently high concentration that it damages the epithelial cell and activates EGFR to induce the production of inflammatory mediators. This possibility is supported by our previous finding that inhibiting the endocytosis of *C. albicans* by treating epithelial cells with cytochalasin D also blocks fungal-induced damage [32].

The central role of EGFR in mediating the host inflammatory response was demonstrated in the immunocompetent mouse model of OPC. Treatment of mice with gefitinib inhibited *C. albicans*-induced phosphorylation of both EphA2 and EGFR in the oral epithelial cells, and reduced the tissue levels of CXCL1/KC, CCL20, S100A8 after both 1 and 2 days of infection. As a result, the accumulation of neutrophils and inflammatory monocytes in the oral tissues was dramatically decreased. In the gefitinib-treated mice, the levels of IL-1α, IL-1β, IL-17A, and TNFα were only reduced at a single time point, suggesting that the production of these cytokines may be governed independently of EGFR.

Inhibition of EGFR has different effects in different models of mucosal infection. Recently, Ho et al. [18] reported that in the zebrafish swimbladder model of mucosal candidiasis, gefitinib treatment impaired the accumulation of neutrophils and led to enhanced mortality, even though it had no effect on fungal burden. Our findings demonstrate that in immunocompetent mice, gefitinib has different effects; although it reduces cytokine production and accumulation of phagocytes, it also decreases oral fungal burden. We and others have found previously that EGFR inhibition also reduces oral fungal burden in steroid-immunosuppressed mice [5, 18]. As inhibition of EGFR reduces *C. albicans* endocytosis by oral epithelial cells *in vitro*, it is probable that the decrease in oral fungal burden by EGFR inhibition is due to impaired epithelial cell invasion *in vivo*.

When we tested the virulence of the *C. albicans* mutants with deletions in *ALS1, ALS3*, and/or *ECE1*, we found that some of the *in vitro* data were recapitulated in mice. After 2 days of infection, mice inoculated with the *ece1*Δ/Δ mutant, the *als1*Δ/Δ *als3*Δ/Δ mutant, and the *als1*Δ/Δ *als3*Δ/Δ *ece1*Δ/Δ mutant all had a similar reduction in oral fungal burden. This finding parallels the reduced epithelial cell damage induced by these strains *in vitro* and indicates that the combination of Als1 and Als3 function along with Ece1/Candidalysin to induce oral infection. It was notable that although the *als1*Δ/Δ *als3*Δ/Δ mutant had extremely impaired adherence to and invasion of oral epithelial cells *In vitro*, its virulence defect in mice was less severe. A likely explanation for this result is that other adhesins and invasins such as Hwp1 [33] and Ssa1 [4] may compensate for the absence of Als1 and Als3 *in vivo*. It is also possible that the *als1*Δ/Δ *als3*Δ/Δ mutant may invade oral epithelial cells *in vivo* by a receptor-independent mechanism, such as by active penetration [2].

The mouse data indicated that different subsets of the host inflammatory response are induced by different combinations of Als1, Als3, and Ece1/Candidalysin. Mice infected with the *ece1*Δ/Δ mutant had reduced levels of CXCL1/KC and CCL20 in their oral tissues, similar to mice that had been treated with gefitinib. Although mice infected with the *als1*Δ/Δ *als3*Δ/Δ *ece1*Δ/Δ mutant had a comparable reduction in these chemokines, animals infected with the *als1*Δ/Δ *als3*Δ/Δ had wild-type levels. Collectively, these results suggest that Ece1/Candidalysin is necessary for *C. albicans* to active EGFR *in vivo*, which stimulates the production of CXCL1/KC and CCL20. These results also indicate that *in vivo*, Als1 and Als3 are dispensable for the induction of this aspect of the inflammatory response.

The data with the mutant strains of *C. albicans* showed that Als1, Als3, and Ece1/Candidalysin stimulate some proinflammatory responses independently of EGFR. While mice treated with gefitinib only had reduced levels of IL-1α and IL-17A at day 1 post infection, mice infected with either the *ece1*Δ/Δ mutant or the *als1*Δ/Δ *als3*Δ/Δ *ece1*Δ/Δ mutant had reduced levels of these cytokines at both days 1 and 2. Thus, at the later time point, Ece1/Candidalysin induces the production of IL-1α and IL-17A independently of EGFR. Also, while the levels of TNFα were not reduced in mice treated with gefitinib, they were consistently decreased only in mice that were infected with the *als1*Δ/Δ *als3*Δ/Δ *ece1*Δ/Δ mutant. These data suggest that the induction of TNFα production is independent of EGFR signaling, but dependent on the combined activity of Als1, Als3, and Ece1/Candidalysin.

The current results indicate that the capacity of Ece1/Candidalysin to induce IL-1β production is dependent on the anatomic site of infection. We found that mice infected with the *ece1*Δ/Δ mutant strains had wild-type levels of IL-1β in their oral tissues, indicating that production of this cytokine is independent of Ece1/Candidalysin. In the mouse model of disseminated candidiasis, Ece1/Candidalysin is dispensable for IL-1β production in the kidneys [34], but essential for IL-1β production in the brains [35]. Thus, *C. albicans* infection induces IL-1β production in the oral mucosa by a different mechanism than in the brain.

A paradoxical finding was that although mice infected with the *ece1*Δ/Δ mutant had a markedly reduced inflammatory response, they also had reduced oral fungal burden. This reduction in oral fungal burden was observed in two strains of mice infected with independent *ece1*Δ/Δ mutants and has also been reported in the immunosuppressed mouse model of OPC [7]. Because of the decreased inflammatory response induced by the *ece1*Δ/Δ mutant, it would have been expected that this strain would be able to proliferate unimpeded in the oral cavity. For example, when mice are infected intravenously with an *ece1*Δ/Δ mutant, there is significantly reduced production of proinflammatory cytokines and accumulation of neutrophils in the brain, and this dampened inflammatory response leads to increased brain fungal burden [35]. One potential explanation for the reduced oral fungal burden in mice infected with the *ece1*Δ/Δ mutant is that even in the absence of Ece1/Candidalysin, there is still some inflammatory response, possibly induced by IL-1β [36] and the residual phagocytes are able to clear the fungus. Alternatively, epithelial cell damage by Ece1/Candidalysin may release nutrients that are vital for fungal proliferation.

Overall, the data presented here indicate that EphA2 and EGFR are mutually dependent for *C. albicans-induced* activation and that the fungal adhesins/invasins Als1 and Als3 function together with Ece1/Candidalysin to cause epithelial cell damage and induce OPC. Although either Als3 or Ece1/Candidalysin is required for *C. albicans* to activate EGFR and stimulate epithelial cells to secrete proinflammatory mediators *in vitro*, induction of much of the oral inflammatory response requires Ece1/Candidalysin-induced activation of EGFR *in vivo*. However, some inflammatory responses are induced independently of both Ece1/Candidalysin-and EGFR. Studies to delineate these additional fungal factors and signaling pathways are currently underway.

## Materials and Methods

### Ethics statement

All animal work was approved by the Institutional Animal Care and Use Committee (IACUC) of the Lundquist Institute at Harbor-UCLA Medical Center. The collection of blood from human volunteers for neutrophil isolation was also approved by the Institutional Review Board of the Lundquist Institute at Harbor-UCLA Medical Center. Informed consent was obtained from all subjects prior to phlebotomy.

### Fungal strains

The *C. albicans* strains used in this study are listed in S1 table. For strain construction, a transient CRISPR-Cas9 system was employed [37]. Primers used for strain construction and confirmation are listed in S2 table. To construct the initial *ece1*Δ/Δ and *als3*Δ/Δ mutants, MH216, a *his1*Δ/Δ NatR derivative of strain SC5314 was utilized [38]. To delete *ECE1*, MH216 was transformed with the Cas9 DNA cassette, ECE1-2 sgRNA DNA cassette, NAT1-5 sgRNA DNA cassette, and *ecelΔ::r1HIS1r1* repair template. *ALS3* was deleted in a similar manner, except that the ALS3-5P sgRNA DNA cassette and *als3Δ::r1HIS1r1* repair template were used. Transformants were selected on complete synthetic medium (CSM) plates without histidine, and screened for nourseothricin sensitivity on YPD + nourseothricin plates. His+ NatS transformants were checked for deletion of *ECE1* or *ALS3* by PCR genotyping using primers ECE1 check up/F and ECE1 check int/R, and primers ALS3 chk up/F and ALS3 chk int/R, respectively. The presence of the *r1HIS1r1* cassette was verified by PCR genotyping using primers ECE1 check up/F and CdHIS1 Check Int/R or primers ALS3 chk up/F T and CdHIS1 Check Int/R. To generate the *als3*Δ/Δ *ece1*Δ/Δ double mutant, the *als3*Δ/Δ mutant MH562 was independently transformed with the Cas9 DNA cassette, the ECE1-2 sgRNA DNA cassette, and the *ece1Δ::r3NAT1r3* repair template. Transformants were selected on YPD plates containing 400 μg/ml nourseothricin and NatR transformants were PCR genotyped to verify deletion of *ECE1* as described above.

The second set of *C. albicans* mutants was constructed in strain SC5314 using the transient CRISPR-Cas9 system [37] combined with the Nat flipper approach [39, 40]. Briefly, the gRNA and Cas9 constructs were amplified from vector pV1093 (S3 Table) by PCR [41]. The gene deletion constructs with the maltose promoter-driven flippase and actin promoter-driven clonNAT selective marker were amplified with primers carrying micro-arms that were homologous to the flanking regions of target genes from a vector derived from pSF2A-mScarlet [39]. The gRNA, Cas9, and repair constructs were mixed and transformed into *C. albicans* cells using the lithium acetate heat-shock method [42]. Transformants were selected by growth on yeast extract peptone dextrose (YPD) agar containing with 200μg/mL of clonNAT. Successfully gene deletion was confirmed with multiple rounds of diagnostic PCRs, including confirming the integration of the deletion construct at the gene locus and the absence of the target gene, gRNA and Cas9 constructs. To recycle the dominant selective drug marker, confirmed transformants were cultured in YPM (YPD in which glucose was replaced with 2% maltose) broth for 2 days in a 30°C shaking incubator. 200-500 cells were then plated onto YPM plates. Individual colonies from YPM plates were then replicate plated onto YPM+clonNAT. NatS colonies were then picked and analyzed by PCR. Chromosomal rearrangement of constructed mutant strains was excluded by diagnostic PCR.

For the experiments, the *C. albicans* cells were grown for 18 h in YPD broth in a shaking incubator at 30°C. The fungal cells were harvested by centrifugation, washed twice with phosphate-buffered saline (PBS), and counted using a hemacytometer.

### Oral epithelial cells

The OKF6/TERT-2 immortalized human oral epithelial cell line was kindly provided by J. Rheinwald (Harvard University, Cambridge, MA) [25] and was cultured as previously described [24]. OKF6/TERT-2 cells were authenticated by RNA-Seq [43] and tested for mycoplasma contamination. The TR146 buccal mucosa squamous cell carcinoma epithelial cell line was generously provided by J. R. Naglik (Kings College London, UK) and was cultured as previously described [18].

### Inhibitor and agonists

The EGFR kinase inhibitor gefitinib (Selleckchem) was dissolved in DMSO and used at a final concentration of 1 μm. It was added to the host cells 60 min prior to infection and remained in the medium for the entire incubation period. Control cells were incubated with a similar concentration of DMSO at a final concentration of 0.1 %. EFNA1-Fc (Acro Biosystems) was used at a final concentration of 1 μg/ml.

### siRNA

To knockdown EGFR and EphA2, OKF6/TERT-2 cells were transfected with siRNA as described previously [24]. Briefly, the cells were grown in 6-well tissue culture plates and transfected with 80 pmol EGFR siRNA (sc-29301, Santa Cruz Biotechnology), and EphA2 siRNA (sc-29304, Santa Cruz Biotechnology), or a similar amount of random control siRNA (sc-37007, Santa Cruz Biotechnology) using Lipofectamine 2000 (Thermo Fisher Scientific) following the manufacturer’s instructions. The extent of protein knockdown was verified 72 h later by immunoblotting with specific antibodies. Knockdown of the protein of interest was > 80% (S1 Fig)

### Immunoblotting

OKF6/TERT-2 cells or TR146 cells in 24-well tissue culture plates were switched to supplement free KSF medium or DMEM/F12 medium, respectively for 1 h and then infected with 1 × 10^6^ *C. albicans* yeast for various times as described previously [16]. Next, the cells were rinsed with cold HBSS containing protease and phosphatase inhibitors, detached from the plate with a cell scraper, and collected by centrifugation. After boiling the cells in sample buffer, the resultant lysate was separated by SDS-PAGE, and phosphorylation was detected by immunoblotting with specific antibodies against pEphA2 (#6347, Cell Signaling) and pEGFR (#2234, Cell Signaling). Next, the blot was stripped, and the total amount of each protein was detected by immunoblotting with antibodies against EphA2 (D4A2, Cell Signaling), EGFR (#4267, Cell Signaling), and β-actin (#a5441, Sigma). Each experiment was performed at least 3 times.

### Immunoprecipitation

OKF6/TERT-2 cells were grown in 75 cm^2^ flasks to confluency, switched to KSF medium without supplements for 3 h, and then infected with 1×10^8^ *C. albicans* yeast. After 30 or 90 min. OKF6/TERT-2 were washed with ice-cold cold PBS (with Mg^2+^, and Ca^2+^), scraped from the flasks, and lysed with 100 μl ice-cold 5.8% octyl β-D-glucopyranoside (0479-5g; VWR) in the present of protease/phosphatase inhibitors. Whole cells lysates were precleared with 20μl of protein A/G plus (sc-2003; Santa Cruz Biotechnology) at 4°C for 30minutes. The bead-protein mix was centrifuged at 3000 rpm for 30 sec at 4°C and supernatants were collected. 2 μg of anti-EGFR antibody (sc-101; Santa Cruz Biotechnology) or anti-EphA2 antibody (#6347, Cell Signaling) was added to 500 μg of protein and incubated on a rotator at 4°C for 2 hours. 25μl of protein A/G plus was added to each immunoprecipitation sample and incubated for an additional hour at 4°C. Samples were pelleted at 3000 rpm for 30 sec, and washed 3 times in 500 μl of ice-cold 1.5% octyl β-D-glucopyranoside. Proteins were eluted with 30 μl of 2X SDS buffer and then heated at 90°C for 5 minutes. Samples were centrifuged at 3000 rpm for 30 sec after which the supernatants were collected, separated by SDS-PAGE, and analyzed as described above.

### Measurement of epithelial cell endocytosis

The endocytosis of *C. albicans* by oral epithelial cells was quantified as described previously [44]. OKF6/TERT-2 or TR146 oral epithelial cells were grown to confluency on fibronectin-coated circular glass coverslips in 24-well tissue culture plates and then infected for 120 min with 2×10^5^ yeast-phase *C. albicans* cells per well, after which they were fixed, stained, and mounted inverted on microscope slides. The coverslips were viewed with an epifluorescence microscope, and the number of endocytosed organisms per high-power field was determined, counting at least 100 organisms per coverslip. Each experiment was performed at least 3 times in triplicate.

### Cytokine and chemokine measurements *in vitro*

Cytokine levels in culture supernatants were determine as previously described [16]. Briefly OKF6/TERT-2 cells or TR146 cells in a 96-well plate were infected with *C. albicans* at a multiplicity of infection of 5. After 8 h of infection, the medium above the cells was collected, clarified by centrifugation and stored in aliquots at −80 °C. The concentration of inflammatory cytokines and chemokines in the medium was determined using the Luminex multipex assay (R&D Systems).

### Epithelial cell damage

The effects of gefitinib on extent of epithelial cell damage caused by *C. albicans* was determined by our previously described ^51^Cr release assay [24]. OKF6/TERT-2 cells and TR146 cells in a 24-well plate were loaded with ^51^Cr overnight. The next day, they were incubated with gefitinib or diluent and then infected with *C. albicans* at a multiplicity of infection of 10. At various time points, the medium above the epithelial cells was collected and the epithelial cells were lysed with RadiacWash (Biodex). The amount of ^51^Cr released into the medium and remaining in the cells was determined with a gamma counter, and the percentage of ^51^Cr released in the infected cells we compared to the release by uninfected epithelial cells. The experiment was performed 3 times in triplicate.

### Mouse model of oropharyngeal candidiasis

Male, 6 week old BALB/c mice were purchased from Taconics and C57BL/6J mice were purchased from Jackson Laboratories. OPC was induced in mice as described previously [24, 45]. Starting on day −2 relative to infection, the mice were randomly assigned to receive gefitinib or no treatment. Gefitinib was administered by adding the drug to the powdered chow diet at a final concentration of 200 parts-per-million. For inoculation, the animals were sedated, and a swab saturated with 2 × 10^7^ *C. albicans* cells was placed sublingually for 75 min. Mice were sacrificed after 1 and 2 days of infection. The tongues were harvested, weighed, homogenized and quantitatively cultured. The researchers were not blinded to the experimental groups because the endpoints (oral fungal burden, cytokine levels, and leukocyte numbers) were an objective measure of disease severity.

### Cytokine and chemokine measurements *in vivo*

To determine the whole tongue cytokine and chemokine protein concentrations, the mice were orally infected with *C. albicans* as above. After 1 and 2 days of infection, the mice were sacrificed, and their tongues were harvested, weighed and homogenized. The homogenates were cleared by centrifugation and the concentration of inflammatory mediators was measured using a multiplex bead array assay (R&D Systems) as previously described [16, 46].

### Flow cytometry

To detect phosphorylation of EphA2, and EGFR in the tongue of *C. albicans* infected mice, the mice were orally infected with *C. albicans* as above. After 1 of infection, the mice were sacrificed, and their tongues were harvested, fixed in 3% PFA for 30 min. After washing a single cell suspension was prepared as described above. Single cells were permeabilized with methanol for 10 min, washed and stained with anti-pEphA2 (#6347, Cell Signaling) or anti-pEGFR (#2234, Cell Signaling) antibodies over night at 4°C. The next day the cells were further stained with anti-rabbit FITC Ab (Abcam, ab6717), CD326 (Ep-CAM)-PE (G8.8, Biolegend), CD45-APC (30-F11; BD Biosciences) and then analyzed with a BD FACSymphony A5 flow cytometer.

The number of phagocytes in the mouse tongues were characterized as described elsewhere [47]. Briefly, mice were orally infected with *C. albicans* as described above. After 1 d of infection, the animals were administered a sublethal anesthetic mix intraperitoneally. The thorax was opened, and a part of the rib cage removed to gain access to the heart. The vena cava was transected and the blood was flushed from the vasculature by slowly injecting 10 mL PBS into the right ventricle. The tongue was harvested and cut into small pieces in 100 μL of ice-cold PBS. 1 mL digestion mix (4.8 mg/ml Collagenase IV; Worthington Biochem, and 200 μg/ml DNase I; Roche Diagnostics, in 1x PBS) was added after which the tissue was incubated at 37°C for 45 min. The resulting tissue suspension was then passed through a 100 μm cell strainer. The single-cell suspensions were incubated with rat anti-mouse CD16/32 (2.4G2; BD Biosciences) for 10 min in FACS buffer at 4°C to block Fc receptors. For staining of surface antigens, cells were incubated with fluorochrome-conjugated (FITC, PE, PE-Cy7, allophycocyanin [APC], APC-eFluor 780,) antibodies against mouse CD45 (30-F11; BD Biosciences), Ly6C (AL-21; BD Biosciences), Ly6G (1A8, BioLegend), CD11b (M1/70; eBioscience), and CD90.2 (30-H12; BioLegend). After washing with FACS buffer, the cell suspension was stained with a LIVE/DEAD fluorescent dye (7-AAD; BD Biosciences) for 10 min. The stained cells were analyzed on a 2-laser LSRII flow cytometer (BD Biosciences), and the data were analyzed using FACS Diva (BD Biosciences) and FlowJo software (Treestar). Only single cells were analyzed, and cell numbers were quantified using PE-conjugated fluorescent counting beads (Spherotech).

### Phagocyte killing assays

The effects of gefitinib on neutrophil killing of *C. albicans* were determined by our previously described method [24]. To study human cells, neutrophils were isolated from the blood of healthy volunteers and incubated with gefitinib or diluent in RPMI 1640 medium plus 10% fetal bovine serum for 1 h at 37°C. Next, the neutrophils were mixed with an equal number of serum-opsonized *C. albicans* cells. After a 3 h incubation, the neutrophils were lysed by sonication, and the number of viable *C. albicans* cells was determined by quantitative culture.

To study bone marrow-derived neutrophils and macrophages (BMDMs), bone marrow cells from *BALB/c* mice (Taconics) were flushed from femurs and tibias using sterile RPMI 1640 medium supplemented with 10% fetal bovine serum (FBS) and 2 mM EDTA onto a 50 ml screw top Falcon tube fitted with a 100 μm filter [48]. Mouse neutrophils were purified from bone marrow cells using negative magnetic bead selection according to the manufacturer’s instructions (MojoSort, BioLegend). These neutrophils had > 90% purity and > 90% viability as determined by flow cytometry. To isolate BMDMs, 6×10^6^ bone marrow cells per 75 cm^2^ were seeded in RPMI 1640 supplemented with 20% FBS, 100 μg/ml streptomycin, 100 U/ml penicillin, 2 mM Glutamine, and 25 ng/ml rHu M-CSF (PeproTech). After 7 days, the BMDMs were treated with gefitinib or the diluent and then incubated with serum-opsonized *C. albicans* cells (multiplicity of infection 1:20)for 3 h. Next, the BMDMs were scraped, lysed by sonication, and the number of viable *C. albicans* cells was determined by quantitative culture.

### Statistics

At least three biological replicates were performed for all *in vitro* experiments unless otherwise indicated. Data were compared by non-parametric Mann-Whitney, or non-parametric one-way Kruskal-Wallis test followed by Dunn’s post-hoc using GraphPad Prism (v. 8) software. P values < 0.05 were considered statistically significant.

## Data Availability

All relevant data are within the manuscript and its Supporting Information files.

## Acknowledgments

This work was supported in part by NIH grant R00DE026856 to Mark Swidergall, R21AI144878 to Aaron P. Mtichell, and R01DE022600 and R01AI124566 to Scott G. Filler. Michael D. Lazarus. was supported by the LA BioMed Summer Fellowship Program. The funders had no role in study design, data collection and analysis, decision to publish, or preparation of the manuscript. We thank the members of the Division of Infectious Diseases at Harbor-UCLA Medical Center for critical suggestions. The content is solely the responsibility of the authors and does not necessarily represent the official views of the National Institutes of Health.

