## Supplemental figures for "Activation of oral epithelial EphA2-EFGR signaling by *Candida albicans* virulence factors"

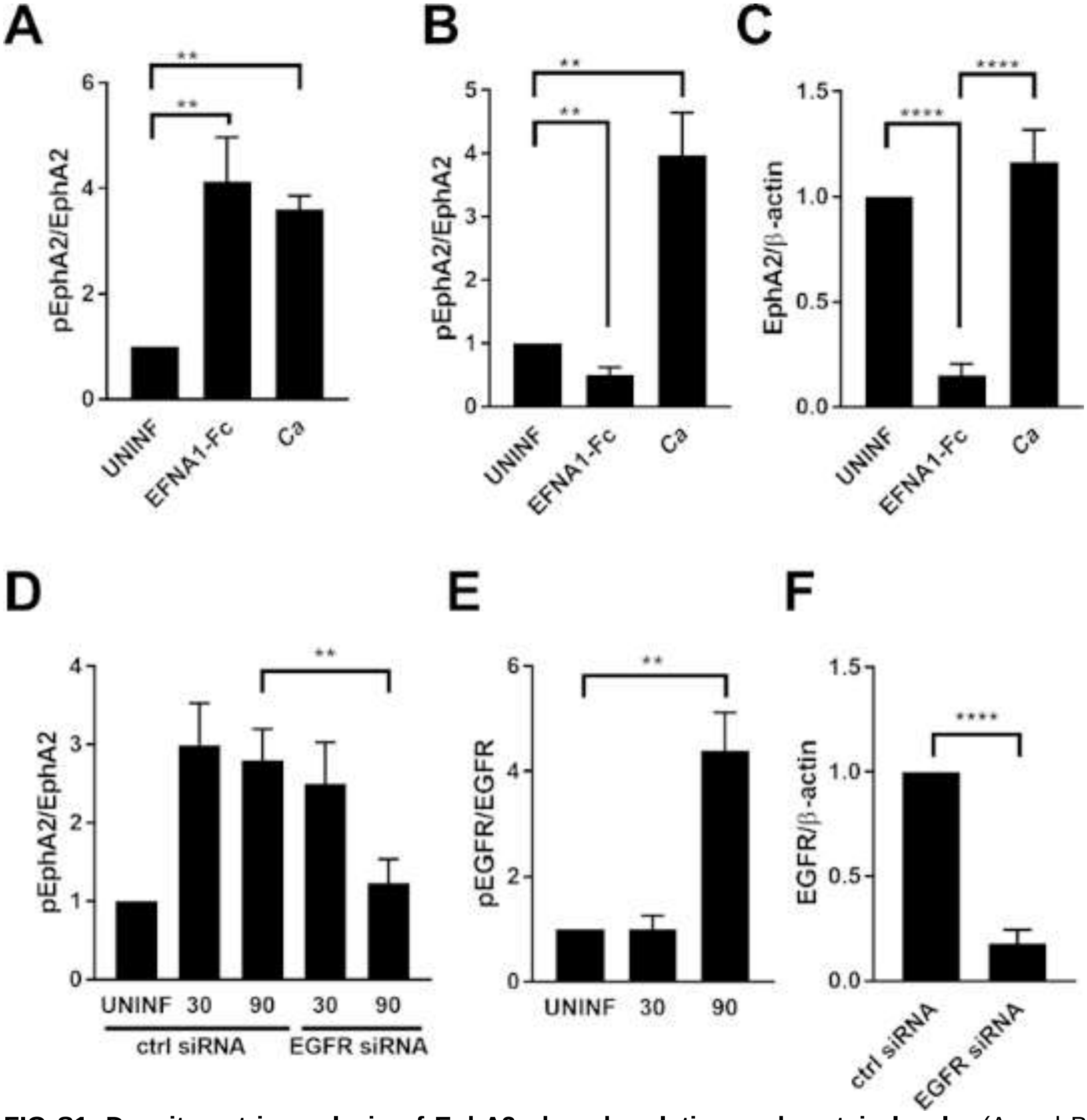


**FIG S1. Densitometric analysis of EphA2 phosphorylation and protein levels.** (A and B). Phosphorylation of EphA2 in uninfected oral epithelial cells (UNINF) and epithelial cells exposed to ephrin A1-Fc (EFNA1-Fc) or yeast-phase *C. albicans* SC5314 (Ca) for 15 min (A) and 60 min (B). Graphs show the relative ratio of phosphorylated EphA2 to total EphA2. (C) Effects of exposure to ephrin A1-Fc or yeast-phase *C. albicans* for 60 min on total EphA2 protein levels. (D) Effects of EGFR siRNA on EphA2 phosphorylation in oral epithelial cells infected with *C. albicans* for the indicated times. (E) Time course of EGFR phosphorylation induced by *C. albicans*. (F) Extent of siRNA knockdown of EGFR. Data are the mean ± SD of 3 independent immunoblots. Images of representative immunoblots are show in Fig. 1. Data were analyzed using the two-tailed Student's t-test assuming unequal variances. **, $p < 0.01$; ***, $p < 0.001$; ****, $p < 0.0001$.

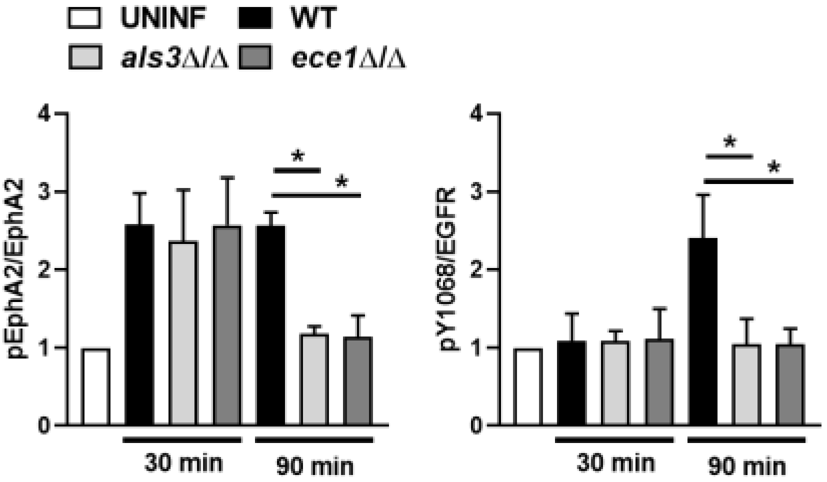


**Fig S2. The *C. albicans als3*Δ/Δ and *ece1*Δ/Δ mutant strains induces weak EGFR phosphorylation and transient phosphorylation of EphA2**. Densitometric analysis of EphA2 and EGFR phosphorylation in oral epithelial cells that had been infected with the indicated *C. albicans* strains of *C. albicans* for 30 and 90 min. Images of representative immunoblots are show in Fig. 2. Data were analyzed using the two-tailed Student's t-test assuming unequal variances. *, $p < 0.05$.

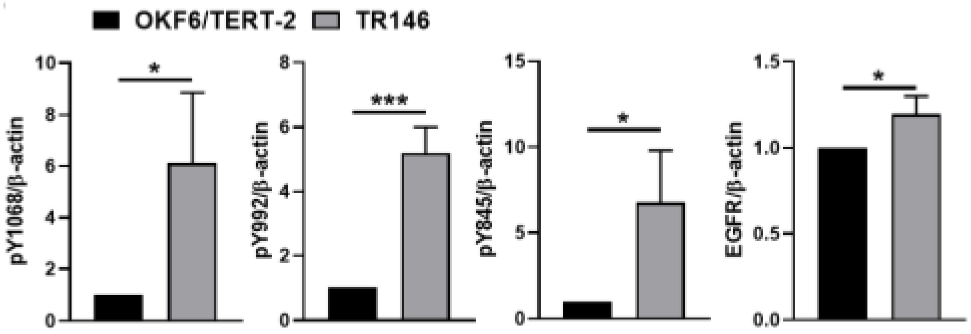


**FIG S3. Densitometric analysis of EGFR phosphorylation and protein levels in unstimulated OKF6/TERT-2 and TR146 oral epithelial cell lines**. Images of representative immunoblots are show in Fig. 4. Data were analyzed using the two-tailed Student's t-test assuming unequal variances. *, $p < 0.05$; ***, $p < 0.001$.

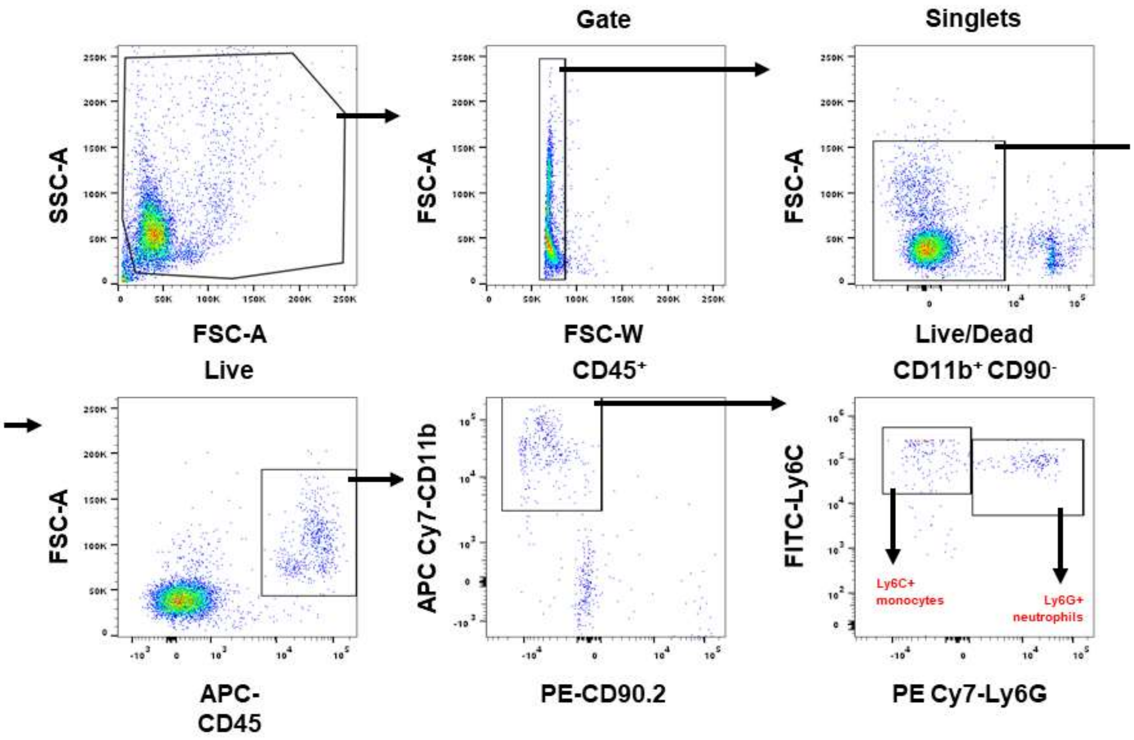


**Fig S4. Gating strategies used to identify Ly6C$^{hi}$ inflammatory monocytes and Ly6C$^{+}$ neutrophils in the flow cytometric analysis of the tongue digests**. The results from these experiments are shown in Fig. 6.

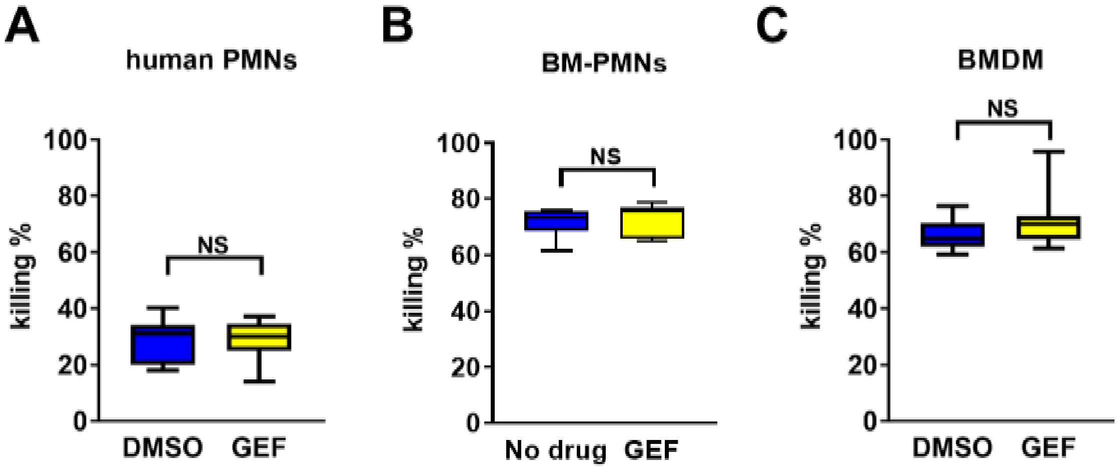


**Fig S5. Gefitinib has no effect on the killing of *C. albicans* by phagocytes**. (A-C) The percentage of *C. albicans* cells killed by human neutrophils (A), bone marrow neutrophils isolated from gefitinib treated mice (B), and mouse bone marrow derived macrophages (BMDM) (B). Data are the combined results of 3 experiments, each performed in duplicate. Data were analyzed using the two-tailed Student's t-test assuming unequal variances. NS, not significant.

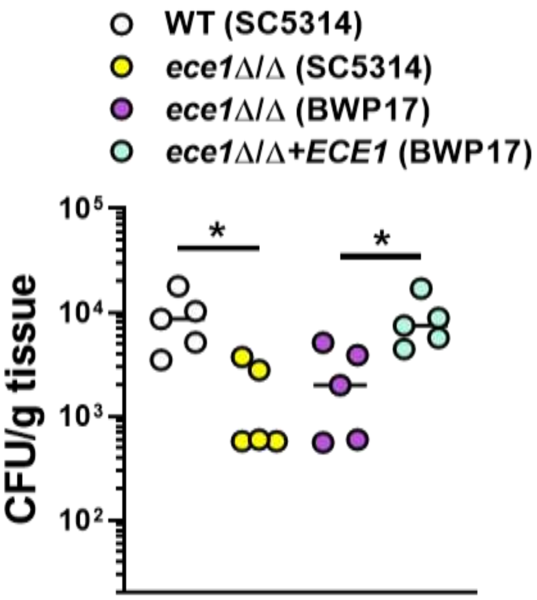


**Fig S6. Deletion of *ECE1* results in lower oral fungal burden.** Oral fungal burden of immunocompetent C57BL/6 mice infected with indicated strains of *C. albicans* after 2 days of infection. Results are median of at total of 5 mice per group from a single experiment. The y-axis is set at the limit of detection (20 CFU/g tissue). Data were analyzed using the Mann-Whitney test.*, $p < 0.05$.

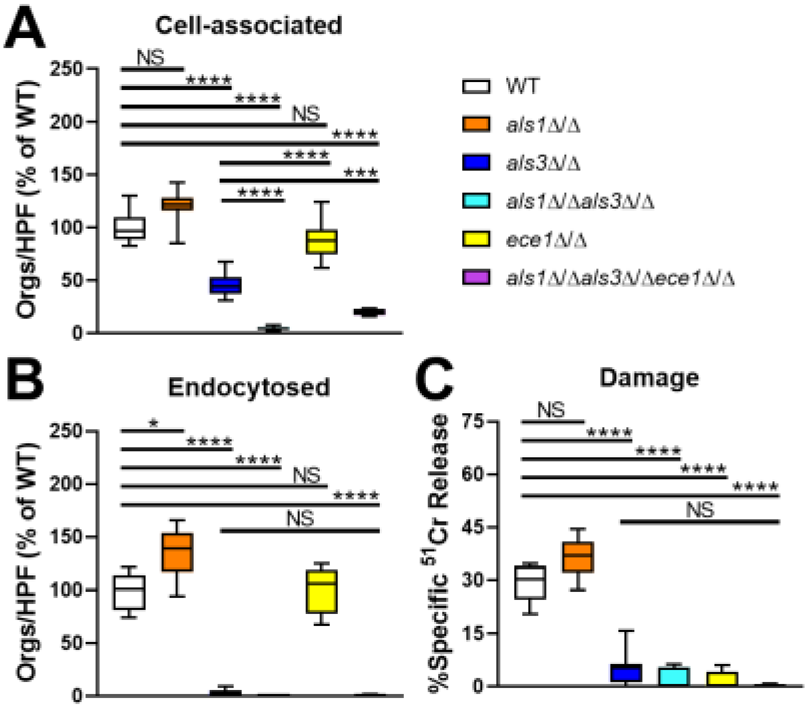


**Fig S7. Epithelial interactions of the indicated *C. albicans* strains.** Data are the combined results of three experiments, each performed in triplicate. Statistical significance was determined by analysis of variance with the Dunnett's correction for multiple comparisons. NS, not significant; orgs/HPF, organisms per high power field; WT, wild-type; *, $p$ <0.05; $p$ < 0.01; ***, $p$ < 0.001; ****, $p$ < 0.0001.
