## Supplemental table 2 for "Activation of oral epithelial EphA2-EFGR signaling by *Candida albicans* virulence factors"

**Table S2. List of primers used in the experments.**

| Name | Sequence |
|---|---|
| sgRNA/F ECE1 | TCATGTTGAATTCTGGAGCAGTTTTAGAGCTAGAAATAGCAAGTTAAA |
| SNR52/R ECE1 | TGCTCCAGAATTCAACATGACAAATTAAAAATAGTTTACGCAAGTC |
| sgRNA/F ECE1-2 | AATTTCTGGCAATCTGACGAGTTTTAGAGCTAGAAATAGCAAGTTAAA |
| SNR52/R ECE1-2 | TCGTCAGATTGCCAGAAATTCAAATTAAAAATAGTTTACGCAAGTC |
| sgRNA/F ALS3-5P | ATTGTTACTCATATATTTGTGTTTTAGAGCTAGAAATAGCAAGTTAAA |
| SNR52/R ALS3-5P | ACAAATATATGAGTAACAATCAAATTAAAAATAGTTTACGCAAGTC |
| sgRNA/F NAT1-5 | GTCACGACGTTGTAAAACGAGTTTTAGAGCTAGAAATAGCAAGTTAAA |
| SNR52/R NAT1-5 | TCGTTTTACAACGTCGTGACCAAATTAAAAATAGTTTACGCAAGTC |
| ECE1 check up/F | CACCCAATAGGATCAGTAAATTCTGC |
| ECE1 check int/R | ATTTGGATTACTTGTGGAATGTTGC |
| ALS3 chk up/F | TATTGGCAACAACATCTTCCGC |
| ALS3 chk int/R | GAGTCAAAGTATTGCTCACAGTACATG |
| NAT1 Check/R | TCAATGGTGGATCAACTGGAACTTC |
| CdHIS1 Check Int/R | GGCGCAACAGATATATTGGTGCTCG |
| ALS3 del rHIS1rSapI/F | CCTCCCTTGAATTGAGGTCTGATAGTTTTTAATTTCATTTTATTATAATTGTATAAACAA CTACCAACTGCTAATATTAGCTCGAGGTCGACGGTATCG |
| ALS3 del rHIS1rKpnI/R | AATTTTTTTTTTGGAGCCAAAAAAACAAAAAACAAACAAATAACAAAAATCTAAAAAGGC GACTATGATGGTATCATCCTCCAATACGCAAACCGCC |
| ECE1 del rHIS1rSapI/F | CAAAATTGTTTTATTTTTGTTTATCTCTACAACAAACAACTTTCCTTTATTTTACTACCAA CTATTTTCCATTCGTTAAACTCGAGGTCGACGGTATCG |
| ECE1 del rHIS1rKpnI/R | TCAGTTACAGCAAAAGTGTCACAAGACTTATGGAATAAAAGATTAAGCTTGTGGAAAA CAAATTTTTATCTGCTGAGCATCCAATACGCAAACCGCC |
| ECE1 del rNATrBamHI/F | CAAAATTGTTTTATTTTTGTTTATCTCTACAACAAACAACTTTCCTTTATTTTACTACCAA CTATTTTCCATTCGTTAAAGCTTTAATGCGGTAGTTTATCACAG |
| ECE1 del rNATrXMAI/R | TCAGTTACAGCAAAAGTGTCACAAGACTTATGGAATAAAAGATTAAGCTTGTGGAAAA CAAATTTTTATCTGCTGAGCATGTGTGGTCGCCATGATCG |
| NAT1 CRIME/R | CACCATGACCTCTATGTTCTGG |
| NAT1 CRIME/F | CAGACGCGTTGAATTGTCC |
| HIS1 CRIME/F | GCGCAAGAAGCCTCAACT |
| HIS1 CRIME/R | GAGCTACAGGGCTTGACC |

| | |
|---|---|
| Cas9-F | ATCTCATTAGATTTGGAACTTGTGGGTT |
| Cas9-R | TTCGAGCGTCCCAAAACCTTCT |
| SNR52_F ar LF | AAGAAAGAAAGAAAACCAGGAGTGAA |
| SNF52 | GCGGCCGCAGTGATTAGACT |
| ENO1_T_ R | GCAGCTCAGTGATTAAGAGTAAAGATGG |
| ENO1_T_ Far-R | ACAAATATTTAAACTCGGGACCTGG |
| gRNA_Sc reenF | GGCTCGAACACAGTACCTCCAGA |
| gRNA_Sc reenR | GGCGGCAAAACTAATTCTTCTCTT |
| CaCas9_S creenF | AATTATCAAAAGACACCTATGACGACG |
| CaCas9_S creenR | TCAACTGTTTCATCACTTTATCGTCAA |
| ALS1_Δ-F | CAATTGAAATGTGAAAGTTTGTTTTTTCGTTTTACTTCATCAGAATTGTTC AAACAACTACCAATTGTTAATATCAGGGTACCGGGCCCCCCCTCGA |
| ALS1_Δ-R | TAATAATAACACGAAGAAAAGATAAATGTGAACTAGATCAAGCCAAAAA GGTGATCATAACAATATAGTCACCGCTCTAGAACTAGTGGATCTG |
| ALS1_gR NA_R | ATGGGGTTCTCCAGTAGTAAGTTTTAGAGCTAGAAATAGCAAGTTAAA |
| ALS1_gR NA_F | TTACTACTGGAGAACCCCATCAAATTAAAAATAGTTTACGCAAGTC |
| ALS1_OR F F | ATGCTTCAACAATTTACATTG |
| ALS1_OR F R | TAGTTACGATTGAGGATTCATTGC |
| ALS3_Δ-F | CCCTTGAATTGAGGTCTGATAGTTTTTAATTTCATTTTATTATAATTGTAT AAACAACTACCAACTGCTAATATTAGGGTACCGGGCCCCCCCTCGA |
| ALS3_Δ-R | GGAGCCAAAAAAACAAAAAACAAACAAATAACAAAAATCTAAAAAGGC GACTATGATGGTATCATCCCCGCTCTAGAACTAGTGGATCTG |
| ALS3_gR NA_R | GTGCACCTTTCACATTAAGAGTTTTAGAGCTAGAAATAGCAAGTTAAA |
| ALS3_gR NA_F | TCTTAATGTGAAAGGTGCACCAAATTAAAAATAGTTTACGCAAGTC |
| ALS3_OR F F | ATGCTACAACAATATACATTGTTAC |
| ALS3_OR F R | GGTAGTGTGATATGGAATATCAAC |
| ECE1_Δ-F | GTTTTATTTTTGTTTATCTCTACAACAAACAACTTTCCTTTATTTTACTAC CAACTATTTTCCATTCGTTAAAGGTACCGGGCCCCCCCTCGA |
| ECE1_Δ- R | CAGCAAAAGTGTCACAAGACTTATGGAATAAAAGATTAAGCTTGTGG AAAACAAATTTTTATCTGCTGAGCATTTCCGCTCTAGAACTAGTGGATCTG |

| | |
|---|---|
| ECE1_gRNA_R | ATTGTTGCTCGTGTTGCCACGTTTTAGAGCTAGAAATAGCAAGTTAAA |
| ECE1_gRNA_F | GTGGCAACACGAGCAACAATCAAATTAAAAATAGTTTACGCAAGTC |
| ECE1_ORF F | ATGAAATTCTCCAAAATTGCC |
| ECE1_ORF R | GCAGATTCAGCTGATCTAG |
