## Supplemental table 3 for "Activation of oral epithelial EphA2-EFGR signaling by *Candida albicans* virulence factors"

**Table S3. List of plasmids**

| Plasmid Name | Description: | Bacterial Marker | Reference |
|---|---|---|---|
| pV1093 | CaCas9/sgRNA expression vector | ampR | [1] |
| pSFS2A-mScarlet | pSFS2A carrying mScarlet, *MAL2* promoter driven flippase and *ACT1* promoter driven *NAT1* | chlR | [2] |
| pMH05 | YEp24 carrying *NAT1* from pNAT at BamHI site | ampR | [3] |
| pMH06 | YEp24 carrying *NAT1* from pNAT at XmaI site | ampR | [3] |
| pMH01 | pRS424 carrying *C.d.HIS1* from pSN52 at KpnI site | ampR | [4] |
| pMH02 | pRS424 carrying *C.d.HIS1* from pSN52 at SapI site | ampR | [4] |


1. Min K, Ichikawa Y, Woolford CA, Mitchell AP. *Candida albicans* gene deletion with a transient CRISPR-Cas9 system. MSphere. 2016;1(3):00130-16.
2. Frazer C, Hernday AD, Bennett RJ. Monitoring Phenotypic Switching in *Candida albicans* and the Use of Next-Gen Fluorescence Reporters. Curr Protoc Microbiol. 2019;53(1):e76. Epub 2019/02/13. doi: 10.1002/cpmc.76. PubMed PMID: 30747494.
3. Huang MY, Woolford CA, May G, McManus CJ, Mitchell AP. Circuit diversification in a biofilm regulatory network. PLoS Pathog. 2019;15(5):e1007787. Epub 2019/05/23. doi: 10.1371/journal.ppat.1007787. PubMed PMID: 31116789; PubMed Central PMCID: PMCPMC6530872.
4. Huang MY, Mitchell AP. Marker Recycling in *Candida albicans* through CRISPR-Cas9-Induced Marker Excision. mSphere. 2017;2(2). Epub 2017/03/21. doi: 10.1128/mSphere.00050-17. PubMed PMID: 28317025; PubMed Central PMCID: PMCPMC5352831.
