## Supplemental table 1 for "Activation of oral epithelial EphA2-EFGR signaling by *Candida albicans* virulence factors"

**Table S1. List of *C. albicans* strains used in the experiments.**

| Strain Number | Markers | Description / Genotype | Citation |
|---|---|---|---|
| SC5314 | | Wild-type | [1] |
| *ece1*Δ/Δ | His+<br>Arg+<br>Ura+ | *ece1*::*HIS1*/*ece1*::*ARG4*;<br>*RPS1*/*rps1*::*URA3* | [2] |
| *ece1*Δ/Δ REV | His+<br>Arg+<br>Ura+ | *ece1*::*HIS1*/*ece1*::*ARG4*;<br>*RPS1*/*rps1*::*URA3-ECE1* | [2] |
| MH216 | NatR,<br>His- | *SC5314*<br>*his1Δ::r3NAT1r3/his1Δ::r3NAT1r3* | [3] |
| MH499 | NatS,<br>His+ | *his1Δ::r3/his1Δ::r3*<br>*ece1Δ::r1HIS1r1/ece1Δ::r1HIS1r1* | This study |
| MH562 | NatS,<br>His+ | *his1Δ::r3/his1Δ::r3*<br>*als3Δ::r1HIS1r1/als3Δ::r1HIS1r1* | This study |
| MH588 | NatR,<br>His+ | *his1Δ::r3/his1Δ::r3*<br>*als3Δ::r1HIS1r1/als3Δ::r1HIS1r1*<br>*ece1Δ::r3NAT1r3/ece1Δ::r3NAT1r3* | This study |
| JL018 | | *ece1Δ/ ece1Δ* | This study |
| JL030 | | *als1Δ/ als1Δ* | This study |
| JL036 | | *als3Δ/ als3Δ* | This study |
| JL050 | | *als1Δ/ als1Δ*<br>*als3Δ/ als3Δ* | This study |
| JL057 | | *ece1Δ/ ece1Δ*<br>*als1Δ/ als1Δ*<br>*als3Δ/ als3Δ* | This study |


1. Fonzi WA, Irwin MY. Isogenic strain construction and gene mapping in *Candida albicans*. Genetics. 1993;134(3):717-28.
2. Moyes DL, Wilson D, Richardson JP, Mogavero S, Tang SX, Wernecke J, et al. Candidalysin is a fungal peptide toxin critical for mucosal infection. Nature. 2016;532(7597):64-8.
3. Huang MY, Woolford CA, May G, McManus CJ, Mitchell AP. Circuit diversification in a biofilm regulatory network. PLoS Pathog. 2019;15(5):e1007787. Epub 2019/05/23. doi: 10.1371/journal.ppat.1007787. PubMed PMID: 31116789; PubMed Central PMCID: PMCPMC6530872.
